# The biophysical basis of protein domain compatibility

**DOI:** 10.1101/2020.12.09.418442

**Authors:** Willow Coyote-Maestas, David Nedrud, Antonio Suma, Yungui He, Kenneth A. Matreyek, Douglas M. Fowler, Vincenzo Carnevale, Chad L. Myers, Daniel Schmidt

**Affiliations:** Department of Biochemistry, Molecular Biology & Biophysics, University of Minnesota, Minneapolis, MN, 55455, USA; Department of Chemistry, Temple University, Philadelphia, PA, 19122, USA; Department of Genetics, Cell Biology & Development, University of Minnesota, Minneapolis, MN, 55455, USA; Department of Pathology, Case Western Reserve University School of Medicine, Cleveland, OH, 44106, USA; Department of Genome Sciences, University of Washington, Seattle, WA, 98115, USA; Department of Bioengineering, University of Washington, Seattle, WA, 98115, USA; Department of Computer Science and Engineering, University of Minnesota, Minneapolis, MN, 55455, USA

## Abstract

Understanding the biophysical mechanisms that govern the combination of protein domains into viable proteins is essential for advancing synthetic biology and biomedical engineering. Here, we use massively parallel genotype/phenotype assays to determine cell surface expression of over 300,000 variants of the inward rectifier K^+^ channel Kir2.1 recombined with hundreds of protein motifs. We use machine learning to derive a quantitative biophysical model and practical rules for domain recombination. Insertional fitness depends on nonlinear interactions between the biophysical properties of inserted motifs and the recipient protein, which adds a new dimension to the rational design of fusion proteins. Insertion maps reveal a generalizable hierarchical organization of Kir2.1 and several other ion channels that balances stability needed for folding and dynamics required for function.

**Summary:** Massively parallel assays reveal interactions between donor domains and recipient proteins govern domain compatibility

## Main text

Protein domains are the basic evolutionary units that allow rapid emergence of new proteins from domain insertion or recombination (*1*). Accordingly, domain recombination-based approaches are often used to generate synthetic proteins in biomedical engineering (*2*). However, synthetically recombined proteins that fold and function well are typically the result of trial-and-error and iterative optimization. Furthermore, deriving practical rules that accelerate domain recombination-based protein design is challenging because structure/function relationships of isolated and recombined domains differ (*3*).

To derive rules for productive domain recombination, we generated 760 polypeptide motif (donor) insertions at all 435 amino acids of the inward rectifier K^+^ channel Kir2.1 (recipient) and then measured cell surface expression of the resulting channel / insertion variants. Previously, we had found surprising variability between three motif’s insertional profiles, which implies complex constraints on donor-recipient compatibility (*4*). We therefore chose 760 donor motifs as a representative sample to exhaustively study compatibility (Supp. Table 1). The massive scale of these experiments (over 300,000 variants) is possible due to insertional libraries with little bias (*5*) (Supp. Fig. 1) and recombining libraries into stable cell lines (*6*).

**Figure 1:**
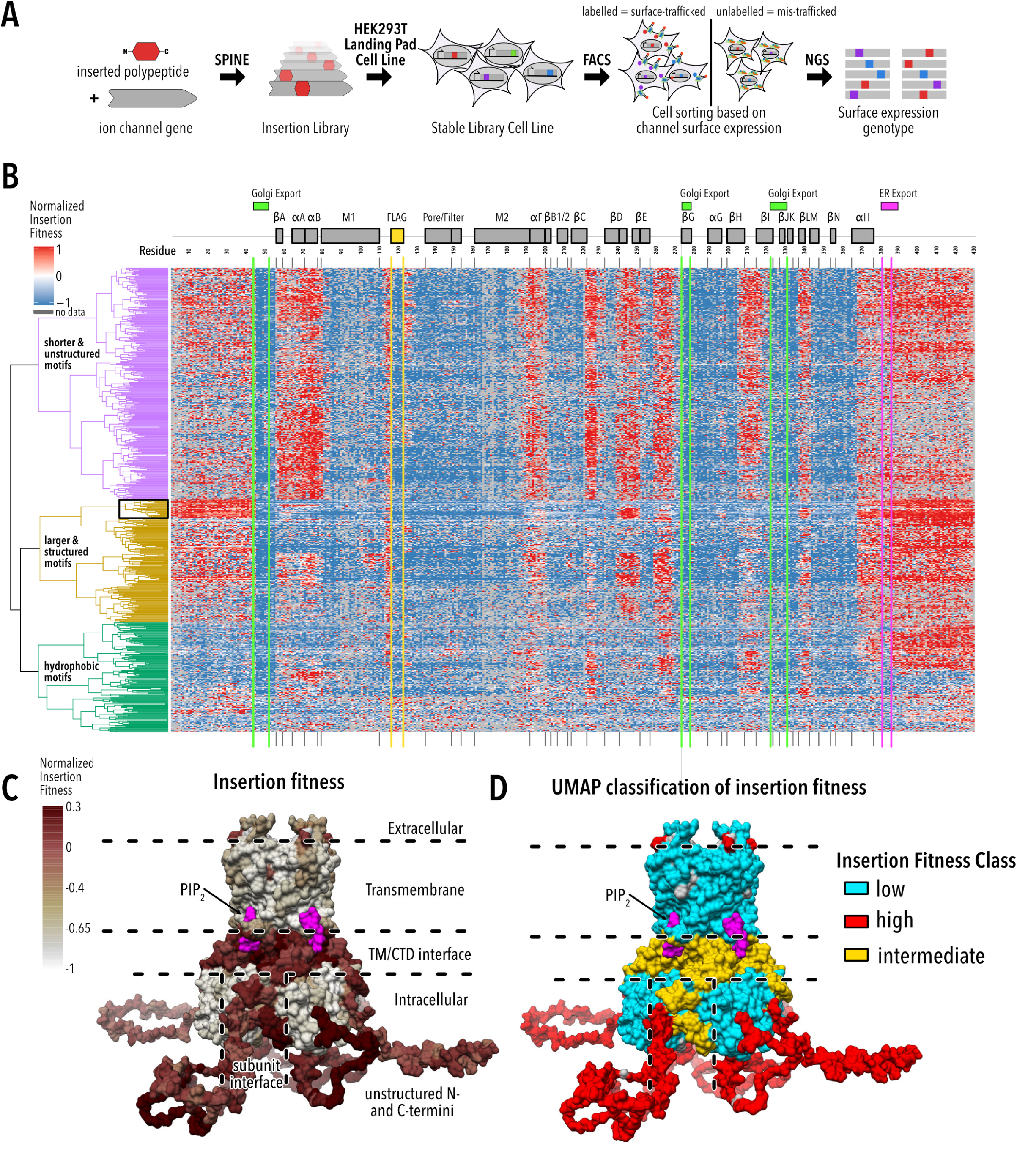
Large-scale insertional fitness profiling. (**A**) Motifs are inserted into all positions of a recipient protein using SPINE (*5*). A stable single-copy insertion library is generated by BxBI-mediated recombination in HEK293T (*6*). Cells are sorted based on channel surface expression determined by antibody labelling of an extracellular FLAG tag. Genotypes of each sorted cell population are recovered by NGS. (**B**) Insertion fitness heatmap of 760 motifs inserted into all positions of Kir2.1. Secondary structural elements (grey boxes) are Kir2.1 are shown above, along known Golgi and ER export signals (green and magenta boxes, respectively). Motifs are hierarchically clustered using a cosine distance metric. Dendrograms are colored by major motifs groups. The black box indicates a subset of ‘well-structured motifs’ (see Fig. 2F-H). **(C-D**) Mean normalized insertion fitness (**C**) or UMAP classification of Kir2.1 insertion fitness (**D**) mapped onto the structure of Kir2.2 (PDB: 3SPI (*21*); 70% identity with Kir2.1; residues 1-40 and 379-410 are modelled). Fitness classes describe conformationally rigid and structured pore domain and CTD beta sheet core (low fitness; cyan), highly flexible and unstructured N/C termini (high fitness; red), and structured yet dynamic interface between TM and CTD, or between subunit in the CTD (intermediate fitness; yellow). PIP_2_ (Kir2.1’s activator) is show in magenta.

### Systematic motif insertions reveal strong fitness pattern consistent with known ion channel biochemistry

For Kir2.1 to maintain cellular excitability (*7*), it must fold, tetramerize, and traffic to the plasma membrane (*8–12*). We measure the impact of insertions on surface expression in Kir2.1 with fluorescent antibody labeling and fluorescently activated cell sorting coupled to sequencing (Fig. 1A). We then calculate surface expression fitness of insertion variants as enrichment or depletion of surface expressed vs. non-surface expressed variants. This data is consistent with expected biochemistry (Fig. 1B-C). Insertions into the extracellular FLAG tag, used to label surface-expressed Kir2.1, mimic decreased fitness because they disrupt antibody binding. Motif insertions into transmembrane regions (M1, M2, Pore, Filter) strongly decrease fitness (Wilcoxon rank sum test p-value < 2.2e-16) presumably by impairing membrane insertion of the nascent protein (*11, 13*). Insertions in folding-critical core beta sheets of the C-terminal domain (CTD) (*14*) also decrease fitness. Conversely, most insertions in the unstructured N- or C-termini are tolerated. As expected, insertions into Golgi export signals decrease surface expression. This is particularly strong for a N-terminal signal with tertiary structure (Fig. 1B, positions 46-50, (*10*)). On the other hand, insertion phenotypes in an ER export signal (the unstructured FCYENE signal (*8*), Fig. 1B, positions 382-387) are more varied with some not affecting surface trafficking. Perhaps the specific residue orientation that is required for function in structured export signal renders them more sensitive to motif insertion, while linear unstructured signals that rely on localized charge or hydrophobicity are more robust. Although insertional fitness patterns are overall consistent with known biochemistry, the variability of insertion fitness across donor motifs and recipient insertion implies more complex mechanisms for domain compatibility.

### Recipient and donor properties interact to determine insertion fitness

To learn if donor properties affect fitness, we hierarchically clustered insertion fitness by motif. This revealed three groups: short unstructured motifs, larger folded motifs, and hydrophobic motifs (Fig. 1B). Unstructured motifs are allowed in many parts of Kir2.1. Structured motifs, which contain nearly all motifs longer than 90 amino acids, are most allowed at the termini and spuriously in structured Kir2.1 regions. Hydrophobic motifs are distinct from other motifs clusters. They decrease fitness in regions (e.g., N terminus) that are universally compatible with the other two motif groups. Some hydrophobic motifs can be inserted where no other motifs can (e.g., beginning of M1 and end of M2 transmembrane helices). Taken together, this suggests that insertion fitness is influenced by the inserted motif’s properties.

To learn if recipient protein properties affect fitness, we used Uniform Manifold Approximation and Projection (UMAP (*15*)) clustering by insertion position. Three distinct clusters emerge (Supp. Fig. 2A) corresponding to contiguous regions of Kir2.1 (Fig. 1D). These regions represent the (1) pore domain and CTD core beta sheets, (2) unstructured N- and C-termini, and (3) PIP_2_ (Kir2.1’s activator) binding sites, interfaces between the pore domain / CTD, and monomer interfaces within CTD. The emergence of discrete contiguous Kir2.1 regions from unbiased clustering suggests that local Kir2.1 properties influence insertional fitness, as well.

**Figure 2:**
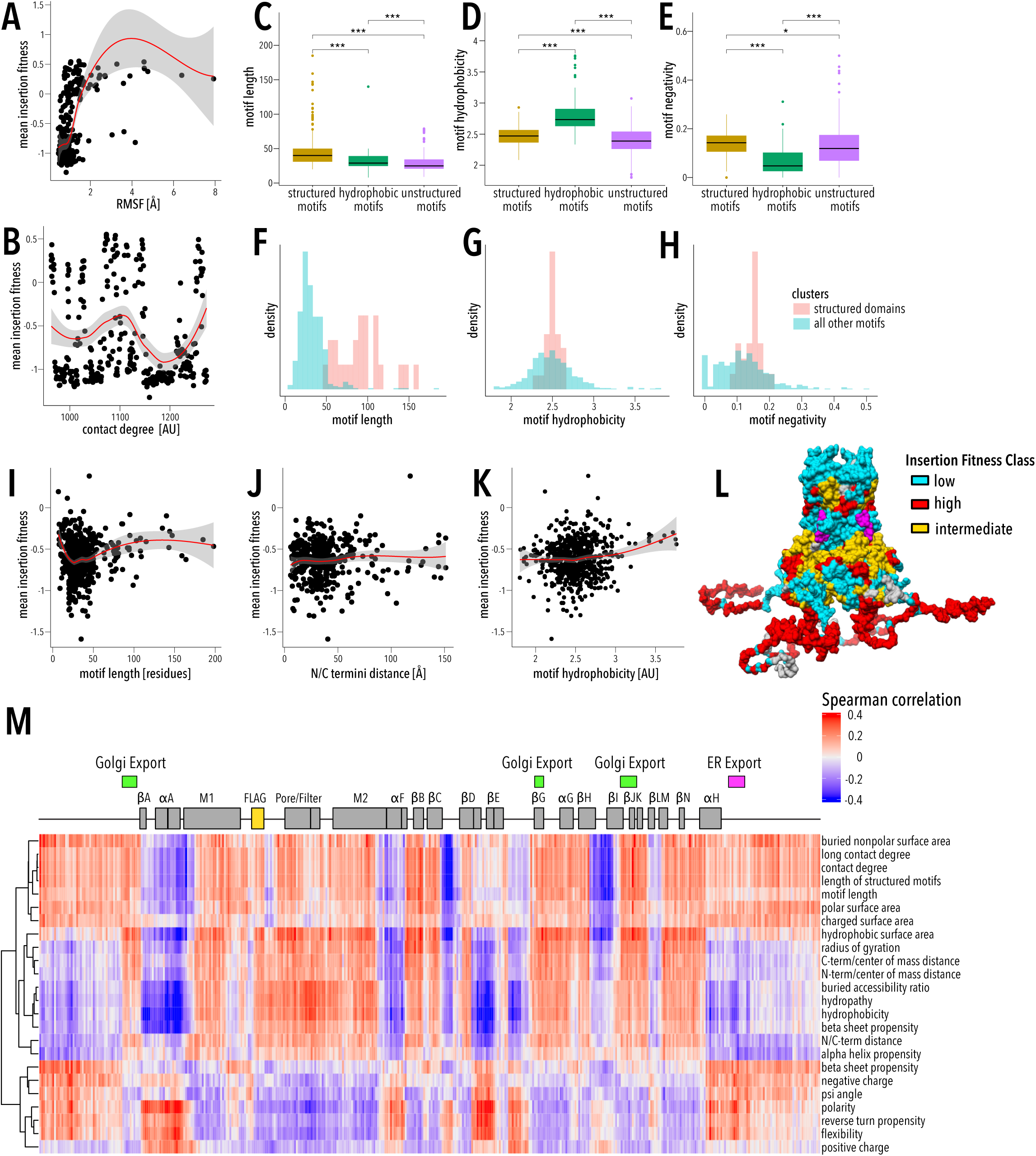
Relationships between fitness data and computed properties. Pairwise scatterplots between recipient properties (**A** – RMSF, **B** – contact degree) and insertion fitness. (**C-E**) Boxplots of motif (**C**) length, (**D**) hydrophobicity, and (**E**) negativity across the three motif clusters from Fig. 1B. Median is marked with a block line, boxes represent the interquartile range, outlier points are shown, and p values from a pairwise Wilcoxon tests are shown. (**F-H**) Density plots of motif (**F**) length, (**G**) hydrophobicity, and (**H**) negativity of the domain cluster and all other motifs colored. Density is weighted group size to allow direct comparison between different sized groups. (**I-K**) Pairwise scatterplots between motif properties (**I** – motif length and **J** – NC termini distance, **K** – motif hydrophobicity) and insertion fitness. (**L**) Hierarchical clusters of motif properties correlations with Kir2.1 position (Supp. Fig. 3) is mapped onto the structure of Kir2.2 (PDB: 3SPI (*21*); 70% identity with Kir2.1; residues 1-40 and 379-410 are modelled). The regulator PIP_2_ is shown in magenta. (**M**) Spearman correlation plot between motif properties and the fitness of that motif at each position. Properties are hierarchically clustered. A LOESS regression curve is fitted to each scatterplot, with the red line represents the fit and the gray area represents the 95% confidence interval. Boxplot significance levels are *** p<0.001, ** p<0.01, and * p<0.05, respectively.

To identify the underlying biophysical properties that influence insertion fitness, we calculated sequence-, structure-, and dynamics-based properties of inserted motifs (Supp. Table 2) and recipient Kir2.1 (Supp. Table 3). We find that insertion fitness has moderately positive correlation with Kir2.1 backbone flexibility (molecular dynamics-derived root mean square fluctuation and anisotropic network model-derived stiffness; Pearson correlation coefficient 0.48 and −0.41, respectively, Fig. 2A) implying that Kir2.1 rearranges structurally after motif insertion. Available space at insertion sites (e.g., contact degree) has a non-monotonic relationship (Fig. 2B). Inserted motif clusters have distinct property distributions. This implies that the pattern of insertion fitness correlates with the biophysical properties of the motif. (Fig. 2C-H). This is illustrated by a subcluster comprised of longer motifs containing hydrophobic and negatively charged residues (black box in Fig 1B, Fig. 2F-H). While motif properties are clearly important, they behave non-linearly. For example, correlation of insertion fitness with motif length is negative for motifs under 25 amino acids but becomes positive for longer motifs (−0.33 and 0.22 Pearson coefficients, respectively, Fig. 2I). Remarkably, all motif properties correlate positively and negatively with fitness dependent on insertion position. Motif lengths, for example, is positively correlated in flexible termini and loops but negatively correlated in the G-loop (Fig. 2M). Our data provide highly resolved information about both donor motifs and the recipient channel that captures the specific rules that govern insertional compatibility (Fig. 2M, Supp. Fig. 3). Hierarchical clustering correlations between fitness and motif properties at each residue separates Kir2.1 into three distinct classes (Fig. 2L, Supp. Fig. 4). These classes are similar to UMAP clustering of fitness alone (compare Fig. 1D and Fig. 2L, Pearson’s χ2 test p-value < 2.2e-16, Cramer’s V 0.42), which indicates that motif and recipient properties can explain insertion fitness. Within each class, correlation sign (positive or negative) between fitness with inserted donor properties is identical. For example, all residues in the pore domain and beta sheet core of the CTD class positively correlate with motif hydrophobicity and negatively with polarity (Supp. Fig. 4). Overall, this suggests that biophysical properties underlie insertional compatibility and properties of Kir2.1 (recipient) and inserted motif (donor) interact to determine fitness.

**Figure 3:**
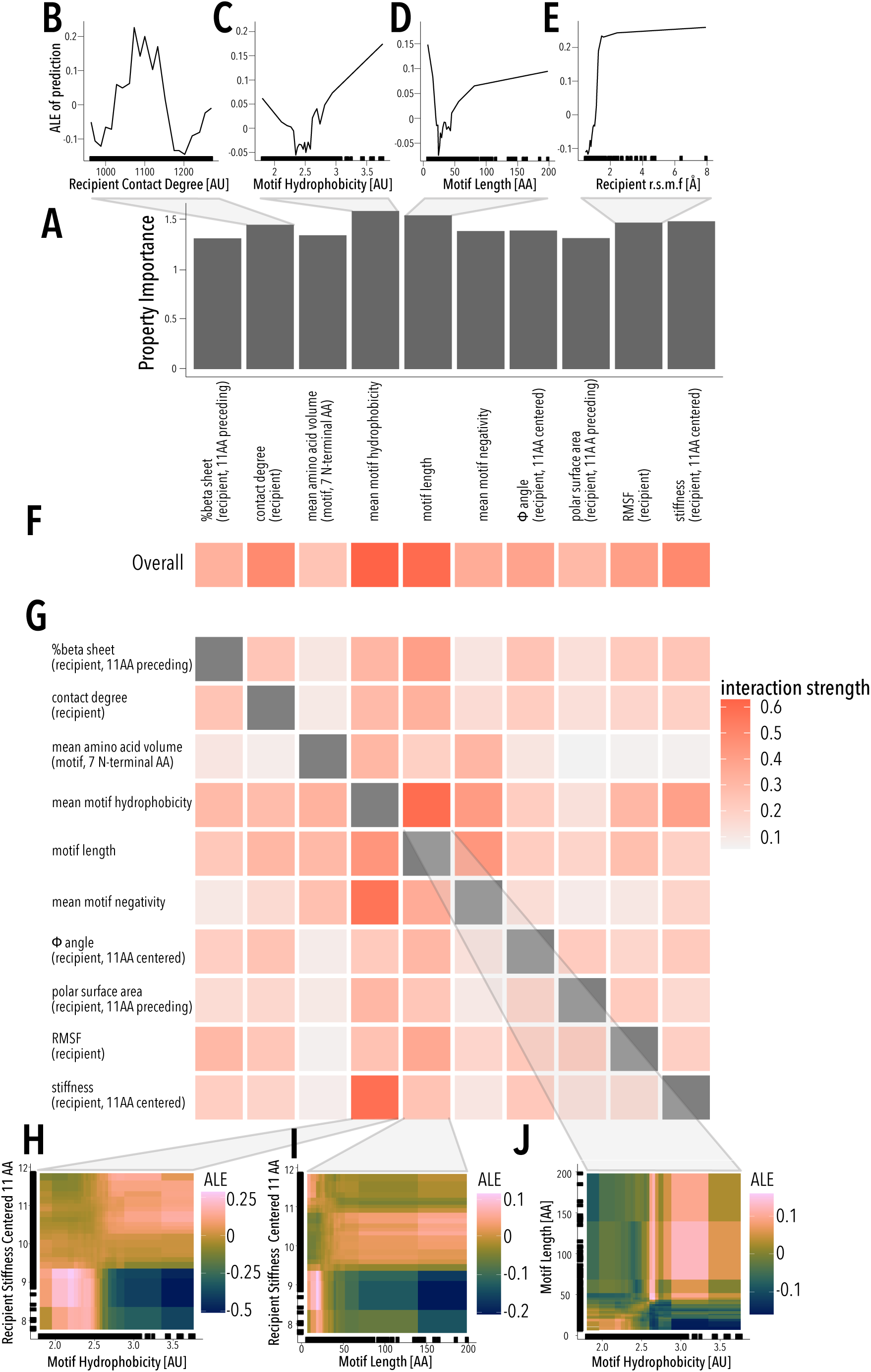
Machine learning model. (**A**) Bar plots of recipient or donor property importance in predicting insertion fitness. Importance is based on the mean absolute error of removing features from the predictive model. (**B-E**) Plots of the Accumulated local effects (ALE) of properties on prediction insertion fitness for (**B**) recipient contact degree, (**C**) motif hydrophobicity, (**D**) motif length, and (**E**) recipient RMSF. (**F, G**) Heatmap of each property’s interaction strength overall (**F**) and pairwise (**G**) with every other property. (**H-J**) Pairwise ALE plots investigate how pairwise interactions contribute to prediction of (**H**) recipient stiffness-motif hydrophobicity, (**I**) recipient stiffness-motif length, and (**J**) motif hydrophobicity-motif length. Pairwise ALE plots are colored from dark blue to pink with increasing ALE scores. Marginal ticks (**B-E, H-J**) indicate values that are covered used in the property data.

**Figure 4:**
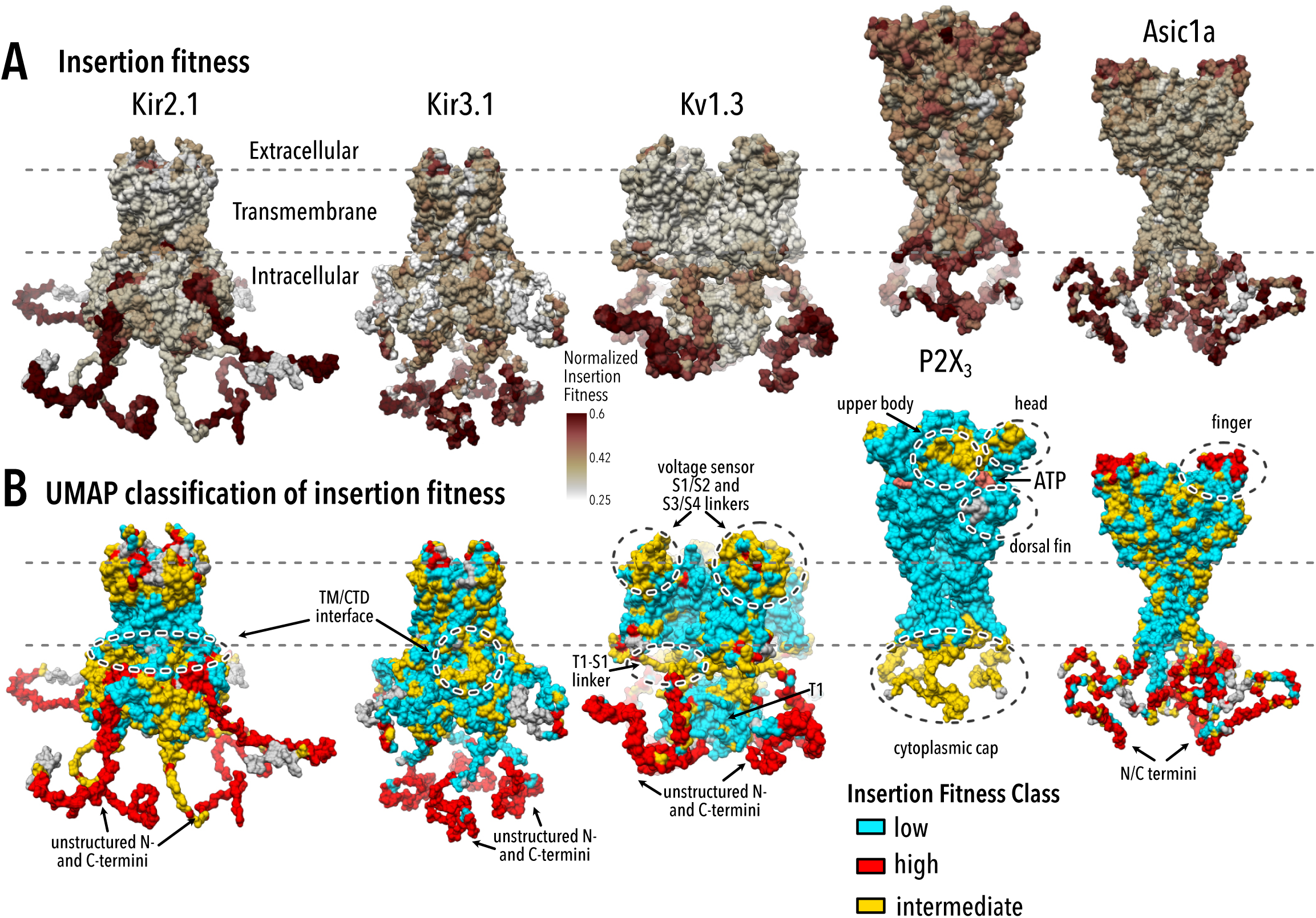
Generalization to other ion channels. Mean insertion fitness (**A**) and UMAP insertion fitness classification (**B**) mapped onto the crystal structures of Kir2.2 (PDB 3SPI (*21*); 70% identity with Kir2.1), Kir3.2 (PDB 4KFM (*23*); 45% identity with Kir3.1), Kv1.2/Kv2.1 paddle chimera (PDB 2R9R (*24*), 62% identical with Kv1.3), P2X_3_ (PDB 5SVK (*25*)), and Asic1a (PDB 6AVE (*26*)). N- and C-terminal residues not resolved in crystal structures are modelled. For all channels apart from P2X_3_, fitness classes describe conformationally rigid and structured regions (low fitness; cyan), highly flexible and unstructured regions (high fitness; red), and structured and dynamic regions (intermediate fitness; yellow) that are often coincide with structural elements important for gating transition (dashed circles). In P2X_3_, there are two regions that separate rigid transmembrane helices and ectodomain (class 1; cyan) and structured and dynamic regions (class 3; yellow). The ligand ATP is shown in soft red.

### Machine learning reveals the basis for donor/recipient compatibility

To identify which donor and recipient properties are important and how they interact in compatible insertions, we used Machine Learning (ML). While ML methods are sometimes treated as black boxes, they are useful for exploring rich genotype/phenotype datasets with non-linear interactions (*16*). We trained and tested regression random forests to predict insertional fitness at every amino acid position based on recipient and motif properties. To identify the most important properties and aid interpretation, we reduced properties from over 900 to 10 based on redundancy and feature importance with little impact on performance (Supp. Fig. 5, Supp. Table 4). The final model successfully predicts insertional fitness for all positions and motifs of data withheld from model training (Supp. Fig. 6).

Local Kir2.1 flexibility (RMSF and stiffness) is important for model performance and is positively associated with insertion fitness (Fig. 3A,E, Supp. Fig. 7C,G). Insertion position space (contact degree) plays a major non-linear role (Fig. 3A,E). Apart from contact degree, all recipient properties have simple monotonic relationships with insertional fitness meaning recipient properties determine whether an insertion is viable (Supp. Fig. 7).

The most important motif properties are length and hydrophobicity, which are both bimodal (Fig. 3C-D). To understand why length and hydrophobicity are bimodal and how properties interact, we explored all property interactions (Fig. 3F-G). Whereas recipient properties do not interact with each other, we find motif properties do interact amongst themselves and with recipient properties (Fig. 3G). This suggests motif property interactions determine insertion fitness and not all insertions are equally compatible with each insertion position.

By exploring property interactions, we learn why different motifs behave distinctly (**Supp. Note 1**, Supp. Figs. 8-10). For example, low contact degree is strongly beneficial for large motifs (Supp. Fig. 8H). Highly hydrophobic short donor motifs are deleterious within flexible regions (small flexible loops) likely because their solvent-exposed hydrophobic residues will be destabilizing and promote aggregation (Fig. 3H-I) (*17*). The small motif cluster contains motifs that are shorter and less hydrophobic, which makes them less disruptive (Fig. 2C-D, Supp. Fig. 9B-C). In contrast, highly hydrophobic motifs are best allowed in buried regions with high stiffness and contact degree because these insertion positions minimize solvent exposure (Fig. 3H-I, Supp. Fig. 10G). Longer motifs benefit from strong positive interactions between motif length and moderate hydrophobicity likely allowing the formation of a hydrophobic core that can promote folding (Fig. 3J, Supp. Fig. 8D) (*18*). Well-folded domains can be stabilizing and promote insertion fitness when there is sufficient space, otherwise large insertions disrupt the recipient protein’s folding (Supp. Fig. 8H). Formation of a stable hydrophobic core as a desirable property of engineered domains corroborates conclusions from high-throughput protein design experiments (*19*).

The ML model allows us to propose practical rules for successfully inserting donor motifs into recipient proteins. Insertion positions are ideally located in flexible protein regions with sufficient space. To form a well-folded domain, motifs need sufficient length and hydrophobic amino acid content to form a well-ordered hydrophobic core. If a desired insertion position is located within a buried and rigid region an inserted motif should be hydrophobic. More flexible regions prefer small non-hydrophobic insertions, and larger more structured domains will only be allowed if there is sufficient space and flexibility. Most significantly, the interactions between motifs and recipient properties determine the outcome of protein recombination. This adds a new dimension to other domain recombination approaches that implicitly treat donor motifs as interchangeable (*20*).

Motif and recipient property interactions produce the distinct classes of motifs and regions (Fig. 1B,D, Fig. 2L). The rigid class with TM and CTD core beta sheets requires specific conformations to achieve a stable fold and allows few insertions. The flexible class with the N/C termini can adopt many conformations and allows most insertions. The class representing interfaces is an intermediate that is structured and dynamic. It contains many Kir2.1 regions (PIP_2_ binding site, TM/CTD and subunit interfaces) that conformationally change upon PIP_2_ binding and during closed to open state transitions (*21, 22*). Since gating mechanisms are conserved across the inward rectifier family (*23*), the interface class may also be enriched for other inward rectifier regulator binding sites, such as Gβγ (GIRK), and ATP (Kir6.2). This is indeed the case (p-value < 2e-16, two-sided Fisher’s Exact test, Supp. Fig. 14). Taken together, distinct class patterns suggest a hierarchical organization of inward rectifiers that balance the stability needed for folding with the conformational dynamics required for function.

### A hierarchical organization of ion channels that balances stability and flexibility for folding and function

To test if our compatibility framework and the hierarchical organization generalizes, we profiled surface expression fitness in the inward rectifier Kir3.1 (GIRK), the voltage-dependent K^+^ channels Kv1.3, the purinoreceptor P2X_3_, and the acid-sensing channel Asic1a by inserting a smaller set of 15 motifs (Fig. 4A, Supp. Table 5, Supp. Fig. 11). Kir3.1 is a G-protein regulated paralog of Kir2.1 with very similar structure (*23*) but requires co-expression of Kir3.2 for effective trafficking (*9*). Kv1.3, P2X_3_, and Asic1a have different folds, gating, and regulation (*24–26*).

The general patterns of surface expression in inward rectifiers also apply to Kv1.3, P2X_3_, and Asic1a. There is weak to moderate correlation between the relative impact of each domain (Supp. Fig. 12-13) in different channels, suggesting that while inserted motifs have similar effects across channels, the recipient channel’s properties dominate. For related channels –Kir2.1 and Kir3.1– insertion profiles are fairly correlated (Pearson correlation coefficient 0.56). Insertions in membrane-embedded regions are deleterious, insertions into termini are allowed, and different inserted motifs give rise to distinct fitness profiles (Supp. Fig. 11). This suggests that properties that dictate fitness in Kir2.1 are generalizable to other ion channels.

Since properties manifested as distinct classes in Kir2.1, we wondered if this concept would also apply to Kir3.1, Kv1.3 and P2X3. Applying the same UMAP-based clustering approach we used for Kir2.1, we find discrete insertion fitness classes in all channels (Fig. 4B). As expected from shared fold architecture, Kir3.1’s classes resemble Kir2.1’s (Pearson’s χ2 test p-value <2.2e-16, Cramer’s V 0.36) with three classes encompassing the TM and CTD core, regulator binding sites and interfaces, and termini. Using established structure/function data, we can infer that classes have distinct roles in folding stability and conformational dynamics. In each channel, there is a class that allows few insertions and corresponds to structural element required for tetramerization (Kv1.3 T1 tetramerization domain), folding (inward rectifier CTD Ig-like fold (*14*), P2X_3_ disulfide-stabilized ecto-domain (*25*), and Asic1a beta sheets), or membrane insertion (transmembrane helices). Most channels have a class that allows nearly all insertions, and which coincides with flexible protein termini. The final class is intermediate, allowing only certain insertions. The intermediate class is enriched for residues that conformationally change during gating or regulation, for example the Kir TM/CTD interface (*21*), the Kv1.3 S1-T1 linker (based on homology of this region to Kv1.2 (*27*)), and P2X_3_ cytoplasmic cap (*25*).

We propose class organization is a universal feature of ion channels that results from constraints on channel structure to satisfy folding, assembly, and interaction with trafficking partners while providing flexibility for allosteric regulation and conformational changes during channel opening and closing. Other studies proposed a similar protein ‘sector’ concept, based on analyzing coevolution of residue pairs in large alignments across homologues (*28*). In contrast, our classes emerge from direct experimental data that are not constrained by statistical modeling’s limitations and reflect underlying biophysical properties. Insertional profiling could be useful as a high-throughput coarse-grain structural biology method to study protein folding and dynamics from steady-state biochemical experiments. Further experiments are required to establish whether the hierarchical organization of insertion fitness extends to all protein classes.

Our dataset provides an unprecedented depth of information across hundreds of inserted donor motifs and several recipient ion channels. Using this dataset, we build a quantitative biophysical model of domain recombination in ion channels. Our discovery of specific interactions between donor and recipient properties is a crucial step towards universal domain recombination ‘grammar’ (*29*) for rational engineering of fusion proteins. Unbiased clustering of insertion fitness reveals a hierarchical organization of ion channels into regions with different material properties (rigid, semi-flexible, flexible) that play distinct roles to balance the stability needed for trafficking and the dynamics required for gating. As a universal organizing framework, this may explain how contradictory requirements for stability and flexibility can be balanced to allow for well-folded and functional proteins.

## Contributions

W.C.-M., D.S., and D.N. conceived the study. W.C.-M. and D.N. generated libraries and performed insertional scans. D.N. coded alignment and enrichment pipelines for data analysis. W.C.-M. carried out machine learning, correlation analysis, and data mining. D.S. conducted clustering analysis and structural mapping. A.S. and V.C. conducted molecular dynamics simulations. K.A.M. and D.M.F. provided reagents and technical advice to construct mammalian cell lines from libraries. C.L.M. provided expertise for random forest model building and data mining. W.C.-M. and D.S. co-wrote the manuscript with input from all co-authors.

## Acknowledgements

We thank James Fraser, Gabriella Estevam, Margaret Titus, the Schmidt Lab, and Fraser lab for helpful feedback and discussion. We acknowledge support from the University of Minnesota Flow Cytometry Resource, in particular Rashi Arora, Therese Martin, and Jason Motl for providing flow cytometry technical support. Andrei Lupas and Vikram Alva kindly provided motif sequences from previous studies. Hellen Farrants and Kai Johnsson kindly provided DHFR and cpDHFR DNA for experiments.

## Supplemental Materials

### Material & Methods

#### Choice of domains

We curated 760 motifs a representative sample of biophysical properties that drive donor/recipient compatibility (Supp. Table 1). Common domains in extant proteins are selected from SMART domain groups, focusing on those with available structural information, and varying range of frequencies within the human genome (*30*). The disordered protein fragments and proteins are from a curated disordered protein database, DISPROT (*31*). The protein fragments are derived from proteins with disordered regions, and the proteins are entire proteins that are disordered. The manually curated motifs include natural, synthetic proteins, several switchable proteins, and a flexible GSAG linker (Supp. Table 5). The polypeptide linkers are manually selected hydrophobic and hydrophilic subsections from Kir2.1. Ancestral motifs have been proposed by Alva et al. (*32*). The small non-domain proteins are manually selected monomeric small proteins which are not commonly recombined. The smotifs are super-secondary structural motifs that are common across proteins (*33*). The natural proteins <50 AA acid motifs are a set of proteins under 50 amino acids that do not contain cysteines that were used in a massive protein stability assay (*19*). Peptide toxins are a set of genetically encodable disulfide-rich neurotoxin peptides.

#### Molecular Biology

Genes encoding human Kir2.1 (Uniprot P63252), human Kir3.1 (Uniprot P48549), human Kir3.2 (Uniprot P48051), human Asic1a (Uniprot P78348), human P2X_3_ (Uniprot P56373), and human Kv1.3 (Uniprot P22001) were produced by DNA synthesis (Twist Bioscience). A Kozak sequence (GCCACC) and P2A-EGFP were added prior and after each open reading frame, respectively. FLAG tag epitopes were added into previous described extracellular loops of Kir2.1 (between S116 and K117 (*12*)), Kir3.1 (between K114 and A115 (*9*)), Asic1a (between F147 and K148 (*34*)), and P2X_3_ (between N72 and R73 based on insertion into paralog P2X_2_ (*35*)). Golden Gate compatible 5’ and 3’ sites were added to each gene by inverse PCR. Sequences of final constructs are in Supplemental Note 2.

#### Library generation

We generated motif insertion libraries using Saturated Programmed Insertional Engineering (SPINE) (*5*). Briefly, we use multi-step Golden Gate cloning to insert a series of motifs in between all consecutive residue pairs of a gene. We break up a gene into fragments (∼169 bp or 53 amino acids) with a genetic handle cassette inserted at every amino acid position. The genetic handle has outward-facing BsaI type IIS restriction sites, which are replaced with any DNA fragment with short N-terminal Ser-Gly and C-terminal Gly-Ser of the inserted motif. We include an antibiotic cassette, chloramphenicol, to remove background wildtype DNA and select for inserted library members. As a quality control step, we sequence all our libraries for baseline coverage prior to screens (Supp. Fig. 20).

#### Cloning domains

The common domains, hand-curated motifs, and non-domain proteins were ordered as gene fragments (Twist Bioscience). The disordered, gene fragments, ancestral, structural, and motifs PDBs <50 amino acids were ordered in the form of an OLS pool (Agilent). All motifs were mammalian codon optimized and designed with amplifiable barcodes and BsaI type IIs restriction sites complementary to those in the inserted genetic handle. Golden gate cloning is conducted with BsaI-v2 HF (NEB), T4 Ligase (NEB) following manufacturer’s instructions. Completed Golden Gate reactions were cleaned with Zymo Clean Concentrate kits and transformed into Lucigen E. cloni™ electrocompetent cells. Diversity was maintained at every step such that there are at least 30x successfully transformed colony forming units as determined by serial dilutions and plating an aliquot of liquid cultures.

#### Library cell line construction

To generate cell lines, we used a rapid single-copy mammalian cell line generation pipeline (*6*). Briefly, insertion libraries are cloned into a staging plasmid with BxBI-compatible *attB* recombination sites using BsaI Golden Gate cloning. We amplify the backbone using inverse PCR and the library of interest with primers that add complementary BsaI cut sites. Golden Gate cloning is conducted with BsaI-v2 HF (NEB), T4 Ligase (NEB) following manufacturer’s instructions. Completed Golden Gate reactions were cleaned with Zymo Clean Concentrate kits and transformed into Lucigen E. cloni™ electrocompetent cells. Diversity was maintained at every step such that there are at least 30x successfully transformed colony forming units as determined by serial dilutions and plating an aliquot of liquid cultures. Completed library landing pad constructs are co-transfected with a BxBI expression construct (pCAG-NLS-Bxb1) into (TetBxB1BFP-iCasp-Blast Clone 12 HEK293T cells). This cell line has a genetically integrated tetracycline induction cassette, followed by a BxBI recombination site, and split rapalog inducible dimerizable Casp-9. Cell are maintained in D10 (DMEM, 10% w/v fetal bovine serum (FBS), 1% w/v sodium pyruvate, and 1% w/v penicillin/streptomycin). Two days after transfection, doxycycline (2 ug/ml, Sigma-Aldrich) is added to induce expression of our genes of interest (successful recombination) or the icasp9 selection system (no recombination). Successful recombination shifts the iCasp-9 out of frame, thus only cells that have undergone recombination survive, while those that haven’t will die from iCasp-9-induced apoptosis. One day after doxycycline induction, AP1903 (10 nM, MedChemExpress) is added to cause dimerization of Casp9 and selectively kill cells without successful recombination. One day after AP1903-Casp9 selection, media is changed back to D10 + Doxycyline (2 ug/ml, Sigma-Aldrich) for recovery. Two days after cells have recovered, cells are reseeded to enable normal cell growth. Once cells reach confluency, library cells are frozen in glycerol stocks in aliquots for assays.

#### Sequencing-based surface expression assay

To measure how inserted motifs disrupt channel expression, we measured surface expression of all variants. We thawed glycerol stocks of library cell lines into wells of a 6 well dish, swapped media the following day to D10, grew cells to confluency, split once to ensure maximum cell health, and swapped media for D10 + doxycycline (2 ug/ml, Sigma-Aldrich). Kir3.1 cannot homo-tetramerize and therefore requires a co-expressed Kir3.2 or Kir3.4 inward rectifier to surface express (*21*). For this reason, 48 hours prior to sorting Kir3.1 libraries, we transiently transfected the stable Kir3.1 insertion library cell line with 2 ug Kir3.2-P2A-miRFP670 and 6ul Turbofect per well of a 6 well plate. For all libraries except for Kv1.3, we detached cells with 1 ml Accutase (Sigma-Aldrich), spun down and washed three times with FACS buffer (2% FBS, 0.1% NaN_3_, 1X PBS), incubated for 1-hour rocking at 4degC with a BV421 anti-flag antibody (BD Bioscience), washed twice with FACS buffers, filtered with cell strainer 5 ml tubes (Falcon), covered with aluminum foil, and kept on ice for transfer to the flow cytometry core. For Kv1.3, cells were detached and washed the same except after initial washing cells were brought up in FACS buffer with Agitoxin-2-Cys-TAMRA (5nM, Alomone), filtered with cell strainer 5 ml tubes, and brought to cell sorting facility on ice. Before sorting, 5% of cells were saved as a control sample for sequencing prior to sorting.

All cells except for Kir3.1 were sorted into unlabeled and labeled (either BV421 or Agitoxin-Cys-TAMRA) populations based on EGFP^high^/label^low^ and EGFP^high^/label^high^, respectively. On a BD FACSAria II P69500132 cell sorter, EGFP fluorescence was excited with a 488 nm laser and recorded with a 525/50 nm bandpass filter and 505 nm long-pass filter. BV421 fluorescence was excited using a 405 nm laser and recorded with a 450/50 nm bandpass filter, TAMRA fluorescence was excited using a 561 nm laser and recorded with a 586/15 nm bandpass filter, and miRFP670 was excited with a 640 nm laser and recorded with 670/30 nm bandpass filter.

All cells (expect those expressing Kir3.1) were gated on forward scattering area and side scattering area to find whole cells, forward scattering width, and height to separate single cells, EGFP for cells that expressed variants without errors (our library generation results in single base pair deletions that will not have EGFP expression because deletions will shift EGFP out of frame (*5*)), and label for surface expressed cells. Kir3.1 library cells were gated on forward scattering area and side scattering area to find whole cells, forward scattering width and height to separate single cells, miRFP670 5 times to get varying levels of Kir3.2 co-expression, GFP for cells that expressed variants without errors, and label for surface expressed cells. For simplicity, we only report Kir3.1 enrichment for one level of Kir3.2 (Kir3.2 #4). The surface expression label gate boundaries were determined based on unlabeled cells from the same population because controls tend to have non-representative distributions. Examples of the gating strategy for each channel is depicted in Supplemental figures 14-18.

EGFP^high^/label^low^ and EGFP^high^/label^high^ cells were collected into catch buffer (20% FBS, 0.1% NaN_3_, 1x PBS. For larger pooled sublibrary samples, we collected between at least 100,000 to 500,00 cells per gate which is 8-35x coverage. 15,000 cells in both gates of a Kir2.1 library with a small flexible ASGASGA linker was collected each day to normalize all the pooled libraries. For smaller 15 motifs samples, we collected between 4,000-50,000 of each sample/library pair which is ∼10-120x coverage for all libraries. We find the more disruptive an insertion the more difficult it is to collect sufficient surface-labeled cells to reach 30x coverage. This means that our lower coverage is assuming all positions are represented in surface expressed cells.

#### Sequencing

DNA from pre-sort control and sorted cells were extracted with Microprep DNA kits (Zymo Research) and triple eluted with water. The elute was diluted such that no more than 1.5ug of DNA is used per PCR reaction and amplified for 20 cycles of PCR using Primestar GXL (Takara Clonetech), run on a 1% agarose gel, and gel purified. Primers that bind outside the recombination site ensure leftover plasmid DNA from the original cell line construction step is not amplified. Purified DNA was quantified using Picogreen DNA quantification. Equal amounts by mass of each domain insertion sample were pooled by cell sorting category and split into two domain sets per channel library set to segregate highly similar motifs sequences. Final amplicon pools were as follows: control, surface expression low 1, surface expression high 1, function low1, function high 1, surface expression low 2, surface expression high 2, function low 2, and function high 2. Pooled amplicons were prepared for sequencing using the Nextera XT sample preparation workflow and sequenced using Illumina Novaseq in 2×150bp mode. Read count statistics are in **Supplemental Table 6**.

#### Enrichment Calculations

Forward and reverse reads were aligned individually using a DIP-seq pipeline (*36*), slightly modified for SPINE compatibility and for updated python packages. If both forward and reverse reads report an insertion, duplicated domain insertion calls are removed to avoid artificially boosting counts. This pipeline results in .csv spreadsheets indicating insertion position, direction, and whether it is in frame.

Surface expression enrichment was calculated by comparing the change in EGFP^high^/label^low^ to EGFP^high^/label^high^. Enrichment calculation was based on Enrich2 software (*37*) and written in R. Only positions with reads in both label^low^ and label^high^ groups were used in enrichment calculations. For each cell group, the percentage of reads at each position was calculated after adding 0.5 to assist positions with very small counts. Enrichment was calculated by taking the natural logarithm of EGFP^high^/label^high^ percentage divided by the EGFP^high^/label^low^ percentage for each position (i).

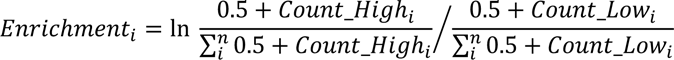

All datasets were z-scored to an internal control flexible linker motif (AGSAGSA) enrichment (separate for each sequencing subpool) by subtracting the average medium enrichment and dividing by the standard deviation of the medium enrichment. Replicates (r) were combined by a weighted average, which was calculated by a restricted maximum likelihood estimate (M) and standard error (SE) using 50 Fisher scoring iterations.

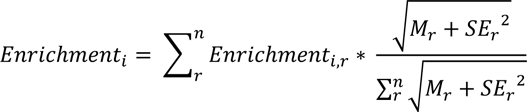

Standard error was calculated assuming a Poisson distribution.

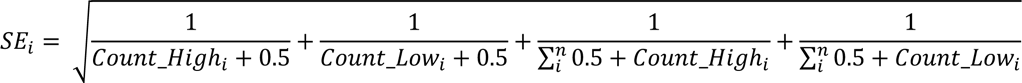

All other positions are treated as NA and are not considered in further analysis (exclusion criteria), except for correlations between datasets as removing data adds more noise than treating NAs as 0s due to sampling.

#### Data quality

Inserting 760 motifs into 435 Kir2.1 positions yields a total theoretical library diversity of 331,360 variants. Each sub-pooled library we generated and screened encompassed 12,500 variants. Due to random variance, some datasets were incomplete (Supp. Fig. 1). To make downstream analysis more robust, we only included motifs with data (after exclusion criteria outlined in *Enrichment Calculations)* in >80% of positions. This left us with 637 out of 760 motifs (further details in Supp. Table 1).

#### Clustering

All motif insertional profiling data was clustered by calculating a cosine distance matrix and clustering it with Ward’s hierarchical clustering method using the hclust function in R with the ‘ward.D2’ method. Uniform Manifold Approximation Projection (UMAP)-based clustering was done using the uwot R package using cosine or Euclidean distance metrics, and a local neighborhood size of 10 sample points. Neighborhood size influences how UMAP balances local versus global structure in the data. Within a range of neighborhood sizes tested (2–50), our choice best conveys the broader structure of the data.

#### Ensemble Network Model

To calculate dynamics of the recipient and motifs with available PDBs, we used the Prody Python package (*38*). For this we used code from from Golinski et al. (*39*) as a starting point kindly provided by Alexander Golinski and Benjamin Hackel (University of Minnesota). We calculated mean stiffness of each backbone based on weighted sums of normal modes from an Anisotropic Network Model of vibration. We calculated summed recipient stiffness for varying lengths (1, 3, 5, 7, 9, 11 amino acids) before, centered on, and after an insertion position. Motif stiffness was summed for the entire motif and for varying lengths of the N- and C-termini (1, 2, 3, 4, 5, and 6 amino acids).

#### Molecular dynamics simulations

All-atom force-field based molecular dynamics simulations were carried out to sample multi-μs trajectories. Our structural models (agonist-bound PDB 3SPI and apo state PDB 3JYC (*21*)) are constituted by the channel embedded in a bilayer of ∼1300 POPC lipids hydrated by two slabs containing ∼170,000 waters and ∼600 KCl ion pairs, for a total of ∼700,000 atoms. We first generated the coordinates of the missing amino acids in the experimental structures (mostly located in unstructured regions) using ROSETTA (for this purpose we generated 10,000 models and kept the representative structure of the most populated cluster). We then used charmm-gui (*40*) to model the bilayer and the aqueous compartment. Simulations are being performed with the charmm36 force field (*41*) at a temperature of T=303.15K, using the highly parallel computational code NAMD2.12 (*42*) on 280 processors cores from Temple University’s Owlsnest. Per-residue root mean squared fluctuations (RMSF) were calculated by considering the position of the C_α_ atoms of each residue using the R bio3D package (*43*).

#### Structure mapping

Calculated properties (e.g., fitness) were mapped onto atomic ion channel structures using Chimera (*44*). Missing loops were manually built using Pymol (*45*) as poly-alanine chains.

#### Amino acid scoring

We calculated bioinformatic scores for amino acids using the Quantiprot python package (*46*). For scores we used: molecular weight, surface area, alpha helical propensity, beta sheet propensity, buried accessibility ratio propensity, flexibility, hydropathy, hydrophobicity, negative charge, pKa, polarity, positive charge, reverse turn propensity, and volume. These scores were calculated for both recipient and donors. We calculated summed recipient scores for varying lengths before, centered on, and after an insertion position (1, 3, 5, 7, 9, 11 amino acids). Motif sequence scores were summed for the entire motif and for varying lengths of the N and C termini (1, 2, 3, 4, 5, and 6 amino acids). Motif length was also included.

#### Protein Structural Properties

A series of properties were calculated with heavily modified code previously used to calculate properties of protein domains kindly provided by Alexander Golinski and Benjamin Hackel (*39*) that uses Pymol (*45*) called from python scripts. Recipient protein PDBs were trimmed of any ions, water, and other non-protein molecules. Recipient protein phi, psi, contact degree, contact order, long contact degree, secondary structure percentage, alpha helical percentage, beta sheet percentage, nonpolar solvent accessible surface area (SASA), charged SASA, and hydrophobic SASA. For each of these properties, we summed recipient structural scores for varying lengths (1, 3, 5, 7, 9, 11 amino acids) before, centered on, and after an insertion position. For motifs with structures, the mean phi angle, mean psi angle, radius of gyration, distance between n and c termini, distance of N and C termini to center of mass, motif size in Daltons, mean contact degree, mean contact order, mean long contact degree, mean secondary structure percentage, mean alpha helical percentage, mean beta sheet percentage, mean nonpolar SASA, mean charged SASA, mean hydrophobic SASA, and RMSD if there were multiple conformers were calculated. In addition to mean motif structural properties, N- and C-terminal varying lengths (1, 2, 3, 4, 5, and 6 amino acids) sums were calculated for the phi angle, psi angle, contact degree, contact order, long contact degree, secondary structure percentage, alpha helical percentage, beta sheet percentage, nonpolar SASA, charged SASA, hydrophobic SASA, and RMSD.

#### Choosing features to train Random Forest

To allow for greater interpretability of our Random Forest-based models, we filtered the input features for redundancy. Our approach to reduce property redundancy was as follows: For motifs, we took the shortest and longest N- and C-terminal features as well as the mean motif features. We identified redundant motif properties by setting a +/− 0.8 correlation cutoff calculated between the motif property and permissibility across all motifs for a given site. We chose the most explanatory of highly correlated motif properties based on summed absolute correlative value across all positions. For recipient properties, we took the longest and shortest of each mean property before, centered and after the insertion position. We identified redundant recipient properties by setting a +/− 0.8 correlation cutoff calculated between the recipient property and permissibility across all positions for a given motif. We chose the most explanatory of highly correlated recipient properties based on summed absolute correlative value across all motifs. These steps reduced our recipient properties from 908 (520 recipient and 388 motif) properties down to 64 (32 recipient and 32 motif) properties.

#### Random Forests

Once we had a non-redundant set of 64 properties, we trained a preliminary random forest model with 500 trees (Supp. Fig. 5). Based on this preliminary model, we further trimmed the properties down to the most explanatory 20 (12 recipient and 8 motif properties). We retrained the model without a significant drop in model performance (39.98% variance explained for 69 properties and 39.44% for 20 properties, Supp. Table 4). However, at this point we were including motif structural properties. This meant that we were not able to include any motifs without structural data. As only 1 of the top 10 most predictive properties (‘Motif Phi Mean’ as the 9^th^ most predictive) were from the structured domain set, we decided to exclude structure-based motif features altogether. This allowed us to include more motifs and reduce our non-redundant properties set down further (39.44% variance explained for 20 properties and 38.69% for 10 properties, Supp. Table 4). We ended up choosing the top 10 most predictive features which included 6 recipient features (stiffness, phi angle of 11 AA centered around insertion site, MD simulation RMSF, contact degree at insertion site, polar surface area of 11 AA preceding insertion site, beta sheet content in 11 AA preceding insertion site) and 4 motif features (mean hydrophobicity, motif length, mean negative charge, mean amino acid volume of 7 N-terminal residues). This final model was trained using 85% of the data, with the other 15% withheld for testing, and performed well on the test dataset (Supp. Fig. 6). All random forests were trained using the Randomforest package in R with 500 trees and localimp = ‘TRUE’ with all model parameters set to default values.

### Supplemental Note 1: Detailed rules for protein recombination from machine learning

#### Properties that guide recombination

Random forest models allow us to study how a set of properties interact non-linearly to give rise to a phenotype. We trained a random forest model on a set of recipient and motif properties to learn what determines productive protein motif insertions into our recipient protein Kir2.1. We calculate feature importance for every property by looking at how model performance is impacted when a given property is not included in the model. We find the most important property overall is motif hydrophobicity, with recipient flexibility (stiffness and RMSF), motif length, and recipient space around an insertion site (contacts) close behind. The most important motif properties are the motifs length and hydrophobicity, and the most important recipient properties are contact degree and stiffness. However, based on feature importance alone, we do not know how properties relate to insertions.

We can further investigate how properties give rise to productive insertions through accumulated local effects (ALE) plots (Figure 3B-E, Supp. Fig. 7). These plots summarize the local effects of a property on the model’s prediction. For example, flexibility appears to have switch-like interactions whereby, below a threshold rigidity, it is quite deleterious (Fig. 3E, Supp. Fig. 7C Positive relationship in RMSF and Supp. Fig. 7G negative relationship in stiffness). Other recipient properties also have straightforward positive or negative relationships such as polar solvent accessible surface area (SASA) (negative, Supp. Fig. 7J), beta sheet % (positive, Supp. fig 7F), and Phi angle (positive, Supp. Fig. 7D). Contact degree on the other hand has a nonlinear and non-monotonic interaction suggesting this recipient property is more complex (Fig. 3B, Supp. Fig. 7E). Overall, recipient features appear to determine insertional fitness in relatively simple ways, such as flexibility and beta sheets 11 amino acids prior to an insertion position are positive, which likely means flexible loops are desirable insertion positions. This result is in line with previous insertion strategies (*20*).

In contrast, all motif properties have more complex relationships to insertional fitness. For example, lower motifs hydrophobicity appears to be deleterious (1.8-2.5) then becomes beneficial at higher values. Similarly, motif length is negative until it becomes beneficial in the model at about 25 amino acids. This is true for the other motif features as well: motif negativity (Supp. Fig. 7B) is initially negative (albeit noisy) then becomes positive. N-terminal 7 amino acid volume (Supp. Fig. 7I) that is initially positive, becomes negative, and returns to be positive. Overall, this suggests motif properties have more complex relationships to insertional fitness. Motif properties are beneficial in some contexts and deleterious in others.

Taken together, recipient properties behave as expected in which flexible loops appear to be beneficial. In contrast to existing approaches to engineer synthetic fusion protein (e.g., (*20*)) that consider inserts to be interchangeable and solely focus on the properties of insertion positions, we propose that inclusion of motifs properties and their interactions is crucial to understand whether an insertion is viable at a given insertion position.

#### Interactions between properties

Random forests are comprised of many decision trees built from random subsets of features that in aggregate predict a desired outcome from properties. Decision trees make predictions by splitting a dataset at property thresholds set on each input feature. Thresholds on multiple input features enable decision trees, and by extension forests, to capture non-linear interactions between properties if they are predictive of the class being modeled. These non-linear interactions are why a property such as motif length can be positive and negative in different contexts.

To interpret why motif properties and contact density behave non-linearly, we explored their interactions (Fig. 3F-J, Supp. Figs. 8-10). We find that motif hydrophobicity and length interact substantially more than all other properties with recipient contact degree and stiffness the next highest interactions (Fig. 3F). Motif properties having more interactions than recipient properties makes sense in light of our earlier observation that motif properties are more likely to non-linearly impact insertional fitness. However, just looking at overall interactions’ strength does not tell us which features interact.

To identify which properties are interacting with which, we calculated pairwise interaction strengths between all properties (Fig. 3G). The strongest interactions overall in-order of strength are pairwise interactions between motif hydrophobicity with motif length, negativity, and stiffness. Overall, there are many pairwise interactions between all the motif features and limited interactions between motif and recipient properties. For recipient properties there are no strong interactions between recipient properties and very few interactions with motif features. That said, recipient stiffness interacts with motif hydrophobicity and length. There are also moderate pairwise interactions between recipient contact degree with hydrophobicity and motif length.

Overall, this means that motif properties interact with each other to determine how a motif behaves when inserted into a position and secondarily with recipient properties to determine whether a motif feature set is beneficial.

To learn which interactions are driving insertional fitness, we calculated and plotted pairwise ALE. It is important to note that pairwise ALE only represents the interaction that contributes to insertional fitness and does not consider how either property contributes alone.

When looking at the strongest interaction overall, motif hydrophobicity and recipient stiffness it is apparent that very high hydrophobicity is extremely deleterious within very flexible regions, low hydrophobicity is very beneficial in flexible regions, and high hydrophobicity is moderately beneficial in stiff (likely buried) regions (Fig. 3H). Observing non-linear interactions help us build hypotheses of underlying biophysical mechanisms, such as hydrophobic residues when exposed and inserted into flexible surface exposed regions are extremely deleterious, whereas when these same motifs are inserted into buried likely more hydrophobic regions these become beneficial. In addition, interactions between motif length (> ∼25 AA) and stiffness demonstrate a different trend, where long insertions into very flexible regions are deleterious (these are regions at the termini of the structure likely needed for folding and small flexible loops) and very rigid regions are also deleterious for long motifs (Fig. 3I). Whereas longer motifs are beneficial in intermediate flexibility regions which are regions within the structured C-terminal domains that move (e.g., flexible loops and the PIP_2_ binding sites). By comparing these two pairwise ALEs (Fig. 3H Stiffness-Hydrophobicity and Fig. 3I Stiffness-Length), we can see that short non-hydrophobics are most preferred within very flexible regions, short hydrophobics are most preferred within very flexible regions, and longer partially hydrophobic motifs are preferred in semi-flexible regions. Furthermore, hydrophobicity is deleterious for short motifs, beneficial for longer motifs, and extremely deleterious for short motifs (Fig. 3J). Perhaps in longer motifs, hydrophobic residues provide stabilization by virtue of well-formed hydrophobic cores, whereas shorter motifs lack well-formed hydrophobic cores and instead expose hydrophobic residues thus becoming very disruptive by promoting aggregation. Overall, this analysis points to motif hydrophobicity and length interacting to determine how a motif behaves within the context of a recipient property. These interactions give rise to the classes of motifs and regions, we observe in clustering (Fig. 1B**, D).**

To further investigate what drove specific motif cluster behavior, we calculated and annotated ALE plots based on where motif class properties are located (Fig. 2C-E).

#### Unstructured short cluster behavior

For the short unstructured motifs, non-hydrophobicity and length are important within unstructured regions because these regions prefer polar hydrophilic motifs as these will be solvent exposed (Supp. Fig. 9B-D). These motifs however are not allowed well in buried regions based on high contacts being deleterious for small motifs (Supp. Fig. 9B, G). In general negativity appears to play a weak negative role (Supp. Fig. 9E). Finally, there is a strong beneficial interaction in regions with beta sheets in the 11 amino acids preceding – perhaps implying flexible loops (Supp. Fig. 9I, J). Flexible motifs are overwhelmingly inserted within flexible loops or at the termini of beta sheets (Supp. Fig. 9A). This class is primarily best allowed within flexible and non-buried regions. Motifs fall into this class if they are non-hydrophobic and small meaning they will be non-disruptive from the perspective of space (contact degree), flexibility (stiffness), and surface exposure (beta sheet %).

#### Hydrophobic motifs

For the hydrophobic motifs, it is quite clear that hydrophobicity drives the behavior of this class. The motif length is not as important because hydrophobic motifs range in size. Hydrophobic motifs mostly benefit from little negativity, which makes sense as many hydrophobic motifs are best allowed with small segments of the transmembrane M1 and negativity would be disruptive when interacting with lipids (Supp. Fig. 10F). Hydrophobic motifs are very deleterious when inserted within very flexible regions and beneficial within rigid regions (Supp. Fig. 10B). This combined with highly hydrophobic motifs being beneficial within high contact regions (Supp. Fig. 10G) means hydrophobics are beneficial when inserted within buried regions. Hydrophobics are highly deleterious in and around beta sheets (Supp. Fig. 10I). Overall, this means hydrophobics behave inversely to the unstructured short cluster. Hydrophobics are mostly deleterious but can be inserted in some buried and transmembrane regions where they will not be disruptive. That said, several recipient flexible loops can accept either motif class (βC-βD, βE-βG, βH-βI, βL-βM). Interestingly, the βD-βE loop and unstructured termini that strongly allows and prefers longer more structured motifs does not allow for most hydrophobic inserts, perhaps because hydrophobics would interact with the solvent to cause misfolding and aggregation.

#### Larger structured motifs

Larger more structured motifs contain nearly all folded proteins and are most interesting from an engineering perspective. This class is overwhelmingly determined by length, with hydrophobicity being intermediate and negativity only slightly higher than other groups. While the overall class does appear to be driven by length, length interacts strongly with hydrophobicity and weakly with negativity (Supp. Fig. 8D-E). Hydrophobicity is positive for long motifs likely representing the ability to form a hydrophobic core and fold. This interaction becomes even more clear when focusing on a subset of motifs within this class that are commonly recombined domains and other well folded larger proteins (Supp. Fig. 8A,D). There is a clear demarcation above which hydrophobicity is highly beneficial (Supp. Fig. 8D), which is likely why folded proteins has such a tight band of hydrophobicity (Fig. 2G). There is a similarly tight distribution of negativity and may be an impact, but it is not nearly as strong (Supp. Fig. 8E-F). Large motifs in very flexible (and generally small loops) large insertions are deleterious but intermediate stiff regions are more amenable to larger insertions (Supp. Fig. 8C). That space is a fundamental determinant for larger motifs is best illustrated by the interactions with contact degree, where low contact degree is beneficial for the largest motifs (Supp. Fig. 8H). Insertions of long motifs appear very deleterious in beta sheet rich regions, which likely disrupt formation of the immunoglobulin-like C-terminal domain of Kir2.1 (Supp. Fig. 8J). Overall, motif length and hydrophobicity strongly interact positively to give rise to increased insertional fitness likely through improving folding. Whether this is beneficial is dependent on where an insertion occurs. Regions with some flexibility and sufficient space are deleterious. However, if there is sufficient space (N -and C-termini and βD-βE loop) insertions are actually quite beneficial. To better design domains for recombination, it would be ideal to have stable domains that have sufficient size and hydrophobicity to be able to maintain their fold after recombination, otherwise their folding thermodynamics will likely be overruled by the recipient protein.

### Supplemental note 2: Channel construct sequences

*Channel sequences:* All channel expression constructs were cloned directly downstream of a Kozak Sequence (GCCACC) and upstream of a P2A-EGFP. Unless otherwise noted, all were cloned into Landing Pad based staging plasmids ‘attB-mcherry’ from the Fowler and Matreyek labs where the mCherry was replaced with our genes of interest. Flag tag is bolded.

Kir2.1_Flag

ATGGGatcAGTGCGAACCAATCGGTATTCTATCGTATCAAGCGAGGAGGACGGGATGAAACTGGCAACCATGGCTGTAGCCAACGGTTTTGGGAATGGGAAATCTAAGGTGCATACCCGCCAACAGTGCCGGTCACGCTTTGTCAAGAAAGATGGCCACTGTAATGTGCAGTTCATAAACGTGGGGGAAAAGGGTCAACGGTACTTGGCCGATATTTTCACAACCTGCGTTGATATCCGCTGGCGCTGGATGTTGGTTATTTTTTGTCTCGCCTTCGTACTGTCTTGGCTCTTCTTTGGCTGCG TTTTTTGGTTGATCGCTTTGTTGCATGGAGATTTGGACACC**GATTATAAAGATGATGATGA TAAA**TCCAAGGTATCCAAGGCCTGTGTCTCCGAAGTAAATTCCTTTACCGCAGCTTTCCTTT TCTCTATCGAAACACAAACCACTATCGGATACGGGTTCCGATGCGTCACAGACGAATGCCCAATAGCCGTTTTCATGGTTGTCTTTCAATCAATAGTGGGCTGTATTATCGATGCATTTATCATTGGGGCCGTGATGGCAAAAATGGCTAAGCCCAAAAAAAGAAATGAGACATTGGTTTTCAGTCACAACGCTGTGATTGCCATGAGGGATGGCAAGCTGTGCCTCATGTGGAGGGTGGGCAATCTGAGAAAGTCCCACCTCGTAGAGGCCCATGTACGAGCACAACTGCTGAAATCACGCATAACTTCAGAAGGAGAGTACATACCACTCGATCAGATTGATATCAATGTGGGCTTCGATAGCGGCATTGACAGGATCTTTCTCGTTAGCCCAATCACCATCGTCCACGAGATTGATGAGGATTCTCCTCTTTATGACCTCTCCAAACAAGACATCGACAACGCTGATTTCGAAATTGTCGTTATACTGGAAGGGATGGTAGAGGCCACCGCTATGACAACCCAATGTCGAAGTAGTTATCTTGCCAATGAAATCCTTTGGGGCCATCGCTACGAACCTGTCTTGTTTGAGGAGAAGCACTATTATAAGGTCGACTACTCTAGGTTCCACAAAACATACGAAGTTCCTAACACACCATTGTGTAGTGCTCGGGACCTTGCAGAGAAGAAGTACATTCTGTCTAACGCAAACTCTTTCTGTTACGAGAACGAAGTAGCTCTCACATCAAAGGAAGAAGAAGAAGATTCAGAGAACGGAGTTCCCGAGTCAACCAGCACCGACTCTCCTCCCGGCATTGACCTGCACAACCAAGCCAGTGTCCCCCTGGAGCCTAGACCTCTTCGGAGAGAGAGTGAAATCGCTGCTTCATCTGCAGTCAATGGGTCAGGC

Kir3.1_Flag

>ATGTCTGCACTTAGGCGGAAGTTCGGTGACGACTATCAGGTCGTGACCACATCCAGTTCA GGATCTGGCCTTCAGCCACAAGGGCCTGGTCAAGGGCCACAACAACAGTTGGTGCCAAAGAAGAAAAGACAGAGGTTCGTTGATAAGAACGGACGATGCAATGTTCAGCACGGCAATCTC GGTAGTGAAACGTCACGCTATTTGTCTGATCTCTTCACCACCTTGGTAGACCTGAAATGGA GGTGGAACCTGTTTATATTCATCCTGACTTATACAGTTGCTTGGCTCTTTATGGCTTCTATG TGGTGGGTTATAGCCTATACTAGAGGTGATCTGAATAAAG**ACTATAAGGACGACGATGAC AAA**GCTCATGTAGGGAATTATACTCCTTGCGTCGCTAACGTCTACAATTTTCCTTCTGCCTT TCTGTTCTTCATAGAGACTGAGGCTACCATTGGGTATGGATACAGGTATATAACCGACAAAT GCCCTGAAGGCATTATTTTGTTTCTCTTTCAATCAATTTTGGGGTCTATTGTAGATGCATTCC TGATCGGGTGTATGTTTATCAAAATGTCACAGCCTAAGAAGAGAGCTGAGACTTTGATGTTT TCCGAGCATGCTGTCATTAGTATGAGGGATGGAAAATTGACTCTTATGTTCAGAGTAGGGA ATCTCCGAAATTCACACATGGTCAGTGCCCAAATCCGGTGTAAACTTCTGAAGAGCAGACA GACCCCAGAAGGGGAGTTTTTGCCTCTTGATCAGCTTGAATTGGATGTGGGTTTTTCCACA GGCGCCGATCAGCTCTTCCTTGTAAGCCCACTGACCATTTGCCATGTCATCGACGCTAAAA GTCCCTTTTATGATCTGAGTCAGAGATCTATGCAGACTGAACAGTTTGAAGTTGTGGTGATA TTGGAAGGTATTGTAGAGACTACTGGGATGACATGCCAAGCACGCACCTCTTATACCGAAG ATGAAGTTTTGTGGGGACACCGATTTTTCCCCGTGATCAGTCTTGAAGAGGGCTTTTTCAA AGTCGATTACTCTCAATTTCATGCTACTTTTGAGGTACCCACTCCACCTTACAGTGTTAAAG AACAAGAGGAAATGCTGCTGATGAGCAGCCCCCTTATCGCACCCGCTATCACCAATTCTAA GGAGAGGCATAACTCCGTTGAATGCCTCGATGGACTGGACGACATTTCTACTAAACTTCCA TCAAAGTTGCAGAAGATAACTGGGCGGGAGGATTTTCCTAAGAAATTGCTGAGGATGTCCT CCACAACTAGCGAAAAAGCATATAGTCTTGGTGACCTGCCCATGAAATTGCAAAGGATTTC AAGCGTTCCTGGTAATTCAGAAGAAAAGCTCGTTAGTAAGACTACCAAAATGCTGTCCGAC CCTATGTCTCAAAGTGTTGCAGACTTGCCACCTAAACTTCAAAAGATGGCAGGAGGCCCTA CTAGAATGGAAGGGAATTTGCCAGCCAAGCTGCGCAAAATGAACTCCGACAGATTCACC

ASIC1a_Flag

ATGGAATTGAAAGCCGAAGAGGAAGAGGTCGGAGGTGTTCAACCAGTTTCTATCCAAGCAT TCGCCTCAAGTTCCACTTTGCACGGGTTGGCACACATATTCTCTTACGAACGCCTCAGCCT CAAACGAGCTCTTTGGGCTCTTTGCTTTCTGGGGTCACTTGCTGTTCTTCTTTGCGTTTGTA CCGAAAGGGTTCAGTATTATTTCCATTATCATCATGTTACTAAACTCGACGAAGTCGCCGCA TCACAGTTGACCTTCCCTGCTGTAACCCTTTGCAATCTGAACGAATTTAGATTTAGCCAAGT TTCTAAAAACGATCTCTATCACGCCGGTGAACTTCTCGCCCTCTTGAATAATCGCTATGAGA TTCCCGATACACAAATGGCAGATGAAAAGCAACTGGAGATCCTCCAGGATAAGGCCAACTT TCGGTCCTTC**GATTATAAAGATGATGATGATAAA**AAGCCCAAGCCCTTCAATATGCGAGAGTTTTACGATCGCGCTGGCCATGATATTCGGGATATGCTTCTCTCATGCCACTTCAGGGGG GAGGTTTGTTCCGCAGAGGACTTCAAGGTCGTGTTTACCCGCTACGGCAAGTGTTATACCT TCAACAGCGGTCGCGACGGGCGCCCTCGGCTTAAAACCATGAAAGGCGGCACTGGTAAC GGACTCGAAATCATGCTGGACATCCAACAAGATGAATACCTCCCCGTGTGGGGTGAAACA GATGAAACCAGTTTCGAAGCTGGTATAAAGGTACAAATACATAGCCAAGATGAGCCCCCCT TTATTGACCAACTTGGCTTCGGTGTAGCACCCGGATTCCAGACATTTGTGGCCTGTCAAGA ACAAAGGTTGATATATCTGCCCCCTCCCTGGGGGACCTGTAAGGCCGTAACAATGGACTC CGACCTGGACTTTTTTGACTCCTACTCCATAACAGCTTGTCGAATTGACTGTGAAACTAGAT ATCTTGTCGAGAATTGTAACTGTAGGATGGTTCACATGCCCGGAGATGCCCCATACTGCAC TCCCGAACAATACAAGGAGTGTGCCGACCCTGCACTTGACTTTCTCGTTGAAAAAGATCAG GAGTATTGCGTGTGCGAGATGCCCTGTAATCTTACACGGTACGGTAAGGAACTTAGTATGG TCAAAATTCCAAGTAAAGCCAGTGCAAAATACTTGGCTAAGAAGTTCAACAAAAGCGAGCA GTACATCGGCGAGAACATTTTGGTTCTCGACATATTCTTCGAAGTCCTGAACTACGAAACTA TTGAACAGAAAAAGGCATACGAGATAGCAGGTCTTTTGGGAGACATAGGAGGGCAGATGG GGCTGTTTATAGGGGCTTCTATTCTGACTGTACTCGAACTGTTTGACTACGCTTATGAAGTC ATTAAACACAAGCTGTGTCGCCGCGGGAAATGTCAGAAGGAGGCTAAGAGAAGCTCAGCC GATAAAGGCGTAGCTCTGTCTTTGGATGATGTAAAACGCCATAATCCTTGCGAATCTCTTC GCGGGCACCCAGCCGGCATGACTTACGCCGCAAACATCCTGCCCCATCATCCCGCACGA GGCACCTTCGAAGATTTTACATGTGCTGCCAGCTCTGCTGTGAATGGTTCTGGA

P2X3_flag

ATGAATTGCATAAGTGATTTTTTTACCTACGAAACCACGAAGAGTGTAGTAGTCAAAAGTTG GACGATAGGAATCATAAACCGCGTCGTACAATTGCTGATTATCTCATACTTTGTAGGCTGG GTTTTTCTGCATGAAAAAGCATACCAAGTTAGGGATACGGCCATTGAGTCATCAGTAGTCA CGAAAGTCAAGGGCAGCGGCCTGTACGCTAAC**GATTACAAGGACGACGATGACAAG**AGG GTTATGGATGTGAGTGATTATGTGACGCCCCCACAGGGGACCAGCGTCTTTGTCATAATAA CCAAAATGATAGTGACGGAAAATCAAATGCAGGGCTTCTGTCCCGAGTCCGAGGAGAAATA CCGATGTGTAAGTGACTCCCAGTGCGGGCCTGAACGCCTCCCTGGTGGTGGTATCCTTAC AGGTCGGTGCGTGAATTATTCAAGTGTGCTGCGGACCTGTGAAATCCAGGGATGGTGTCC AACTGAGGTAGACACAGTCGAGACTCCTATCATGATGGAGGCAGAGAACTTCACTATCTTT ATTAAGAACTCTATACGCTTTCCACTGTTTAATTTTGAGAAGGGTAATCTTCTGCCAAACTTG ACCGCACGAGATATGAAAACATGTAGATTTCATCCGGACAAAGATCCTTTCTGCCCAATTCT CCGCGTTGGAGATGTGGTGAAATTTGCTGGCCAGGACTTTGCAAAGCTGGCACGCACGGGAGGTGTGTTGGGCATAAAAATTGGCTGGGTTTGTGACTTGGACAAGGCCTGGGACCAGTG TATTCCCAAATATTCTTTTACAAGGCTGGACTCAGTATCAGAGAAATCAAGCGTGTCACCCG GTTATAACTTTCGGTTCGCTAAATACTACAAGATGGAGAACGGTTCAGAGTATCGCACCTT GCTGAAGGCGTTTGGTATTCGGTTTGATGTGCTCGTTTACGGTAACGCGGGGAAGTTCAAC ATTATACCGACGATCATTAGCTCCGTGGCCGCTTTTACTTCCGTTGGAGTCGGCACTGTTC TTTGCGATATCATCCTTCTGAATTTTTTGAAAGGAGCCGATCAGTACAAGGCGAAAAAGTTC GAGGAAGTCAATGAAACGACGCTGAAAATTGCAGCGTTGACAAACCCTGTTTATCCAAGCG ATCAAACAACAGCGGAGAAACAGTCCACAGACTCTGGCGCATTTTCTATCGGGCAC

Kv1.3

ATGGACGAGCGGCTCAGTCTACTTCGCTCACCACCACCCCCCTCTGCTCGGCATCGGGCC CATCCCCCTCAACGGCCAGCAAGCAGCGGCGGCGCACACACCTTGGTCAATCACGGGTA CGCCGAGCCCGCTGCTGGCAGGGAACTTCCTCCAGACATGACCGTCGTGCCTGGCGACC ATCTTCTTGAACCCGAAGTGGCAGACGGCGGCGGCGCTCCACCCCAGGGGGGCTGTGGA GGCGGAGGATGCGATCGGTATGAGCCATTGCCCCCCTCCCTTCCAGCTGCCGGCGAGCA AGACTGCTGTGGGGAAAGAGTCGTCATAAACATTAGCGGGCTTCGATTCGAGACACAGCT TAAGACACTTTGTCAATTTCCAGAAACTCTTCTTGGGGACCCAAAACGCCGGATGCGGTAT TTCGACCCCCTTAGAAACGAATACTTTTTTGATCGCAATAGGCCCAGTTTCGACGCCATCCT CTATTACTATCAGAGCGGCGGGCGAATCCGCCGACCTGTCAATGTACCTATCGATATCTTT TCCGAAGAGATCAGGTTTTACCAGCTGGGAGAGGAAGCCATGGAGAAATTTCGCGAGGAC GAGGGGTTTCTGAGAGAAGAGGAACGCCCCCTCCCACGAAGGGATTTCCAGCGACAAGTC TGGCTGTTGTTTGAGTACCCAGAGTCCTCAGGGCCCGCTCGAGGGATAGCAATCGTGAGT GTCCTTGTTATTCTGATTAGTATAGTCATCTTTTGTCTTGAAACACTGCCAGAATTTCGCGAT GAGAAGGATTACCCAGCCTCAACTAGCCAGGACTCATTCGAGGCAGCTGGCAATTCAACA AGCGGGAGCCGGGCAGGTGCATCTTCTTTCTCAGATCCTTTTTTTGTTGTAGAAACACTCT GTATCATTTGGTTCAGTTTTGAATTGTTGGTAAGGTTTTTCGCATGTCCCTCCAAGGCAACA TTCTCCCGAAACATTATGAACCTGATTGATATTGTCGCTATAATACCTTACTTCATCACCCTT GGTACTGAGTTGGCAGAGAGGCAAGGCAATGGGCAGCAAGCCATGTCTTTGGCAATCCTC CGGGTCATCCGGCTTGTGAGGGTTTTTAGGATCTTTAAATTGAGTCGGCATTCAAAAGGGC TCCAGATCCTTGGTCAAACTTTGAAGGCTTCTATGAGAGAACTCGGGCTTCTTATATTTTTT CTCTTCATAGGAGTTATTCTGTTTAGCAGTGCTGTGTATTTCGCTGAGGCCGATGATCCTAC ATCTGGCTTTTCATCAATACCTGACGCATTTTGGTGGGCTGTTGTGACCATGACCACCGTT GGTTACGGTGATATGCACCCCGTGACAATTGGCGGTAAAATCGTGGGCAGCCTCTGTGCAATCGCTGGAGTATTGACCATCGCACTCCCAGTTCCCGTTATTGTTTCCAACTTTAATTACTT CTACCACAGAGAAACCGAGGGAGAAGAACAGAGCCAGTATATGCACGTTGGCTCCTGTCA GCATTTGTCATCAAGTGCCGAGGAATTGCGAAAGGCTCGGTCTAACAGCACCCTGTCCAA GAGTGAGTACATGGTTATCGAGGAGGGAGGTATGAATCATAGCGCTTTCCCCCAGACCCC TTTTAAAACTGGCAACTCTACTGCCACATGCACCACCAACAATAATCCAAACTCCTGCGTCA ACATCAAGAAAATATTTACAGACGTG

Kir3.2 was used but with miRFP670 instead of EGFP and in a transient expression backbone from pEGFPN3.

ATGACAATGGCTAAGTTGACCGAAAGTATGACTAACGTGCTTGAGGGGGACTCCATGGATC AGGATGTGGAGTCACCAGTTGCCATTCACCAGCCTAAGCTGCCCAAACAGGCAAGGGACG ACCTCCCTCGACACATATCACGGGATCGCACTAAGAGAAAGATACAAAGGTATGTAAGGAA AGACGGGAAGTGTAATGTCCATCACGGGAACGTGAGGGAGACATATCGATACTTGACTGA TATCTTCACTACACTGGTGGATCTCAAATGGAGGTTCAATCTGCTCATATTTGTCATGGTTT ATACCGTCACCTGGCTTTTTTTCGGTATGATCTGGTGGCTCATAGCATATATACGGGGGGA TATGGACCATATAGAGGACCCATCATGGACTCCTTGCGTTACAAATCTCAACGGCTTTGTC TCCGCCTTTTTGTTCTCAATTGAAACCGAGACTACAATCGGCTATGGGTACAGGGTCATTA CTGACAAGTGTCCCGAAGGTATCATCCTTCTTTTGATACAATCTGTACTCGGCAGTATTGTT AATGCATTCATGGTTGGCTGCATGTTCGTGAAAATATCCCAGCCCAAAAAAAGGGCTGAGA CATTGGTGTTCTCAACTCACGCAGTAATTTCAATGAGAGATGGCAAGCTCTGTCTTATGTTT CGGGTAGGGGACCTTCGCAACAGCCACATCGTAGAGGCTAGCATCCGGGCAAAACTTATT AAAAGTAAACAGACCAGTGAGGGCGAGTTCATCCCCCTGAACCAGAGTGACATCAACGTG GGATATTATACCGGCGACGATCGCCTGTTTCTCGTTTCACCACTTATAATATCTCATGAGAT CAATCAGCAGAGCCCCTTTTGGGAGATTAGCAAAGCCCAACTCCCTAAGGAAGAGCTTGA GATAGTGGTTATATTGGAAGGAATTGTCGAAGCAACAGGGATGACATGTCAGGCACGGTC CAGTTATATTACATCTGAGATCCTCTGGGGGTACCGGTTTACCCCCGTTTTGACCATGGAA GATGGATTTTATGAGGTCGACTATAATAGTTTTCACGAAACATACGAGACAAGCACACCTTC TCTTTCAGCTAAAGAGTTGGCTGAGTTGGCAAACCGGGCAGAAGTCCCACTTTCCTGGAG CGTGTCTAGCAAGCTGAATCAGCACGCTGAGTTGGAAACCGAGGAAGAGGAGAAGAATCC AGAAGAACTGACTGAGCGGAATGGTGATGTGGCAAATCTCGAGAACGAGTCAAAGGTT

>P2A Sequence used

GCAACTAATTTTAGTCTACTGAAACAAGCTGGTGATGTGGAGGAAAATCCAGGACCA

>EGFP Sequence Used

atggtgagcaagggcgaggagctgttcaccggggtggtgcccatcctggtcgagctggacggcgacgtaaacggccacaagttca gcgtgtccggcgagggcgagggcgatgccacctacggcaagctgaccctgaagttcatctgcaccaccggcaagctgcccgtgcc ctggcccaccctcgtgaccaccctgacctacggcgtgcagtgcttcagccgctaccccgaccacatgaagcagcacgacttcttcaa gtccgccatgcccgaaggctacgtccaggagcgcaccatcttcttcaaggacgacggcaactacaagacccgcgccgaggtgaa gttcgagggcgacaccctggtgaaccgcatcgagctgaagggcatcgacttcaaggaggacggcaacatcctggggcacaagct ggagtacaactacaacagccacaacgtctatatcatggccgacaagcagaagaacggcatcaaggtgaacttcaagatccgcca caacatcgaggacggcagcgtgcagctcgccgaccactaccagcagaacacccccatcggcgacggccccgtgctgctgcccga caaccactacctgagcacccagtccgccctgagcaaagaccccaacgagaagcgcgatcacatggtcctgctggagttcgtgacc gccgccgggatcactctcggcatggacgagctgtacaagtaa

>miRFP670

aTGgTAGCAGGTCATGCCTCTGGCAGCCCCGCATTCGGGACCGCCTCTCATTCGAATTGC GAACATGAAGAGATCCACCTCGCCGGCTCGATCCAGCCGCATGGCGCGCTTCTGGTCGTC AGCGAACATGATCATCGCGTCATCCAGGCCAGCGCCAACGCCGCGGAATTTCTGAATCTC GGAAGCGTACTCGGCGTTCCGCTCGCCGAGATCGACGGCGATCTGTTGATCAAGATCCTG CCGCATCTCGATCCCACCGCCGAAGGCATGCCGGTCGCGGTGCGCTGCCGGATCGGCAA TCCCTCTACGGAGTACTGCGGTCTGATGCATCGGCCTCCGGAAGGCGGGCTGATCATCGA ACTCGAACGTGCCGGCCCGTCGATCGATCTGTCAGGCACGCTGGCGCCGGCGCTGGAGC GGATCCGCACGGCGGGTTCACTGCGCGCGCTGTGCGATGACACCGTGCTGCTGTTTCAG CAGTGCACCGGCTACGACCGGGTGATGGTGTATCGTTTCGATGAGCAAGGCCACGGCCT GGTATTCTCCGAGTGCCATGTGCCTGGGCTCGAATCCTATTTCGGCAACCGCTATCCGTC GTCGACTGTCCCGCAGATGGCGCGGCAGCTGTACGTGCGGCAGCGCGTCCGCGTGCTGG TCGACGTCACCTATCAGCCGGTGCCGCTGGAGCCGCGGCTGTCGCCGCTGACCGGGCGC GATCTCGACATGTCGGGCTGCTTCCTGCGCTCGATGTCGCCGTGCCATTTACAATTCCTGA AGGACATGGGCGTGCGCGCCACCCTGGCGGTGTCGCTGGTGGTCGGCGGCAAGCTGTG GGGCCTGGTTGTCTGTCACCATTATCTGCCGCGCTTCATCCGTTTCGAGCTGCGGGCGAT CTGCAAACGGCTCGCCGAAAGGATCGCGACGCGGATCACCGCGCTTGAGAGCTAA

---

**Supplemental Figure 1:**
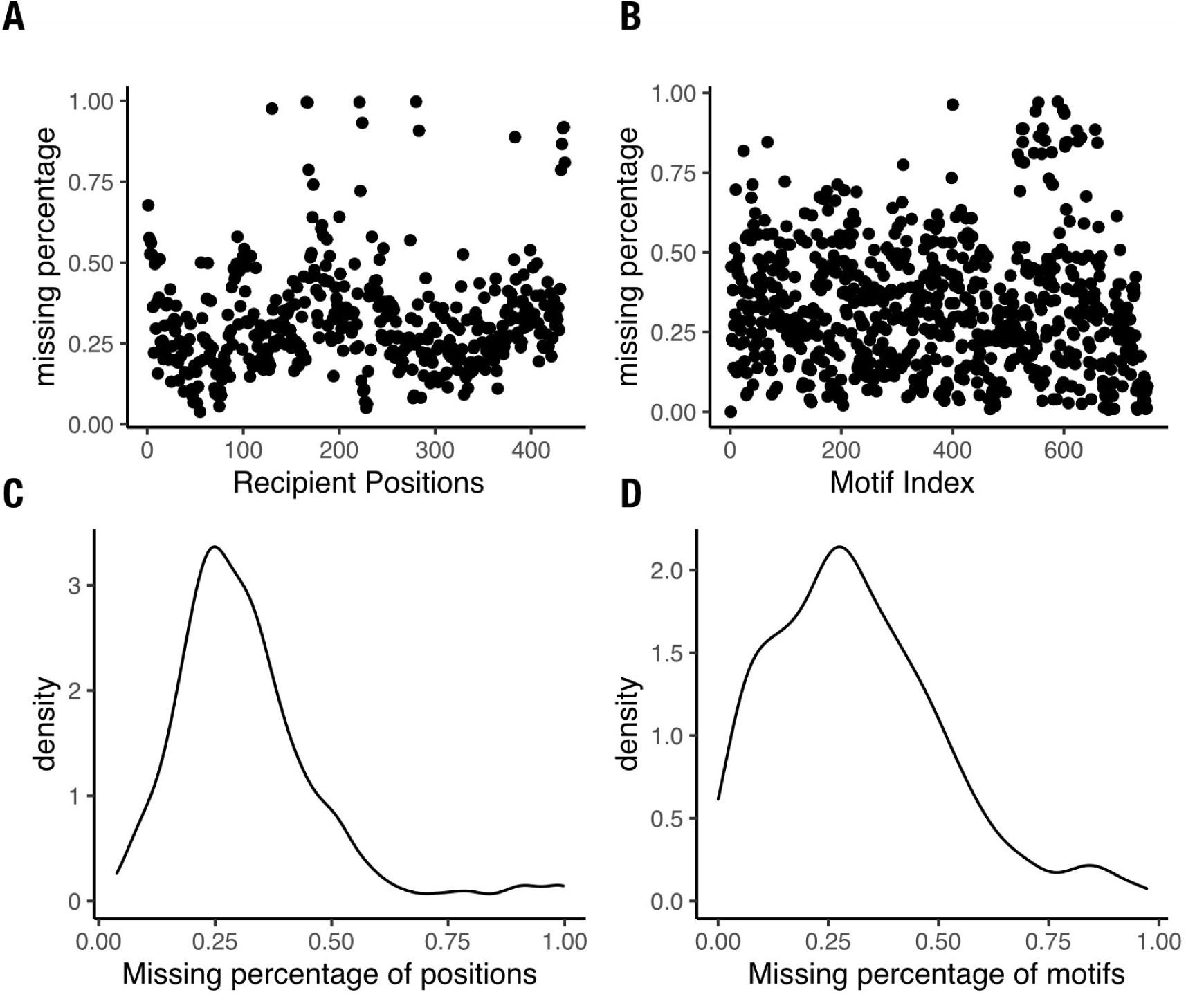
lnsertional fitness coverage. (**A-B**) Scatter plots with the percent missing of Kir2.1 insertion fitness data after alignment by (**A**) position and (**B**) motif. (**C-D**) Density plots of Kir2.1 insertion fitness data percent missing by (**C**) position and (**D**) motif.

**Supplemental Figure 2:**
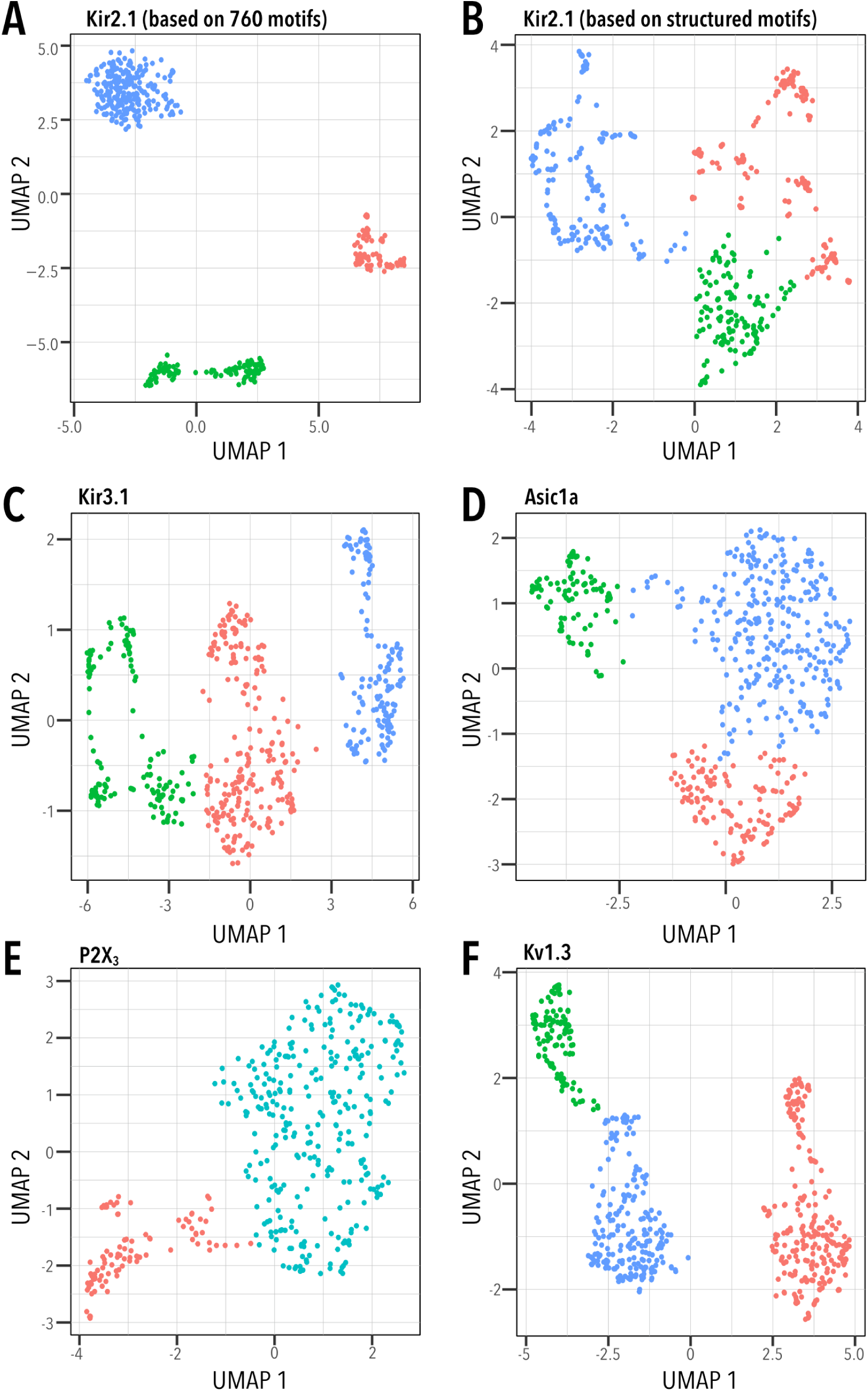
Unbiased clustering of insertion fitness. Uniform Manifold Approximation Projection (UMAP) was used to cluster insertion fitness of each channel. Cluster membership of each residue is indicated by color. Optimal cluster number was determined using Nbclust (47) using the majority rule.

**Supplemental Figure 3:**
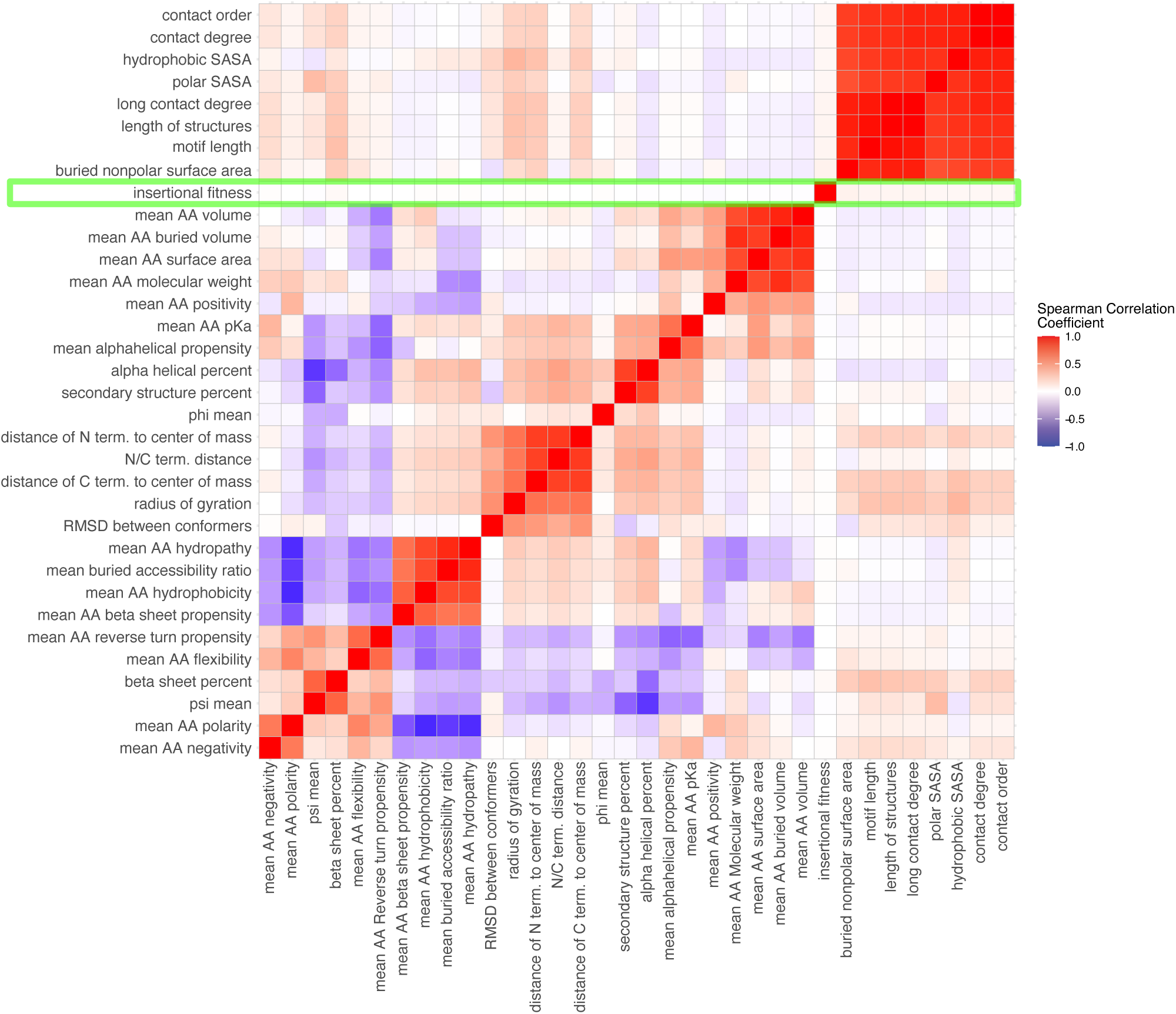
Motif properties and insertional fitness correlations. Correlation plot between motif property and the fitness across all positions. Insertional fitness is not correlated with any motif property. The motif properties and positions are hierarchically clustered (dendrograms not shown) and the plot is colored with spearman correlations increasing from blue-to-red. AA refers to amino acids and SASA refers to solvent accessible surface area.

**Supplemental Figure 4:**
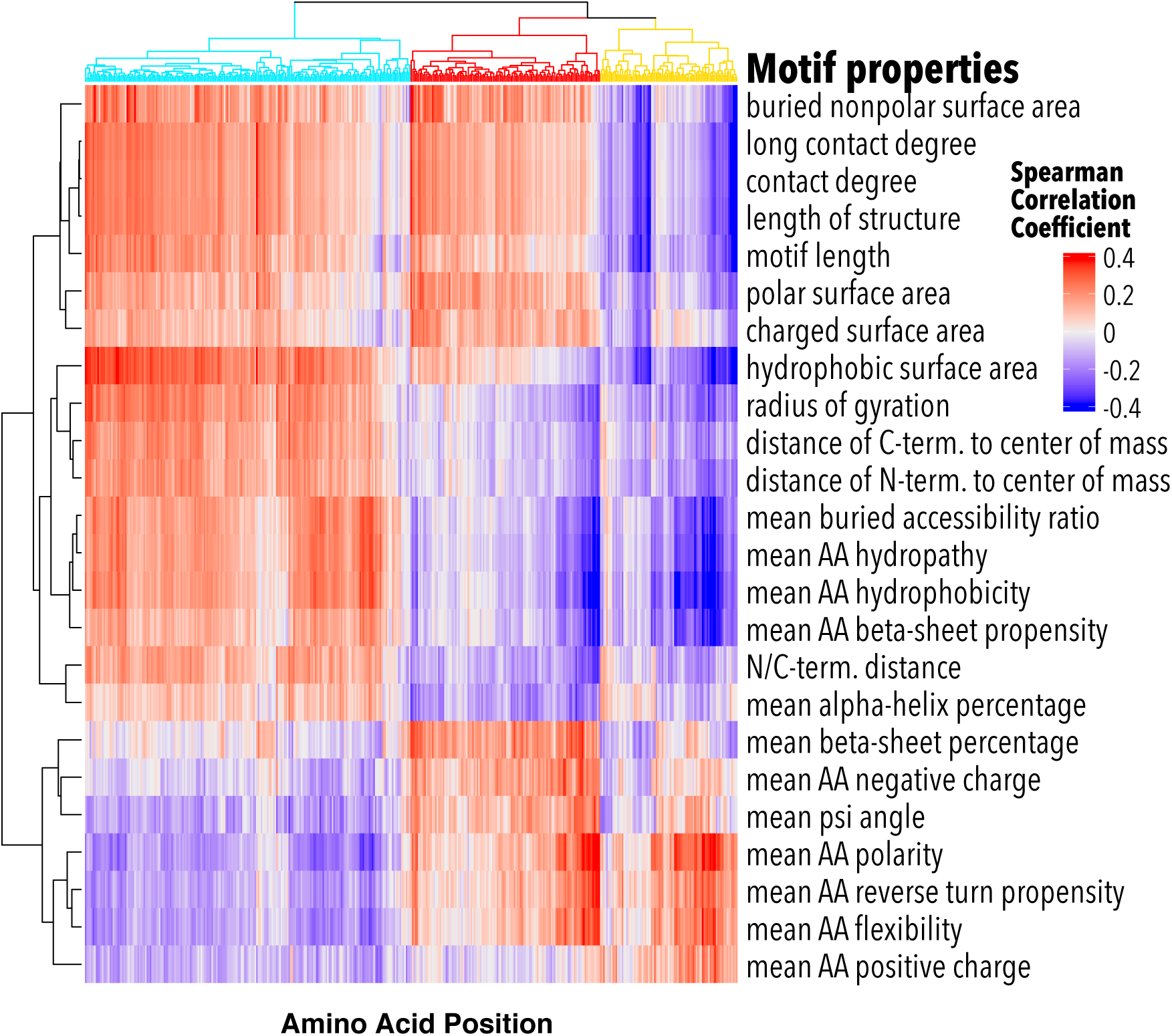
Clustered positions and properties correlation plot. Correlation plot between motif properties and the fitness of that motif at each position. The motif properties and positions are hierarchically clustered. Position clusters dendogram branches are colored (cyan, red, yellow) as in Fig. 2L.

**Supplemental Figure 5:**
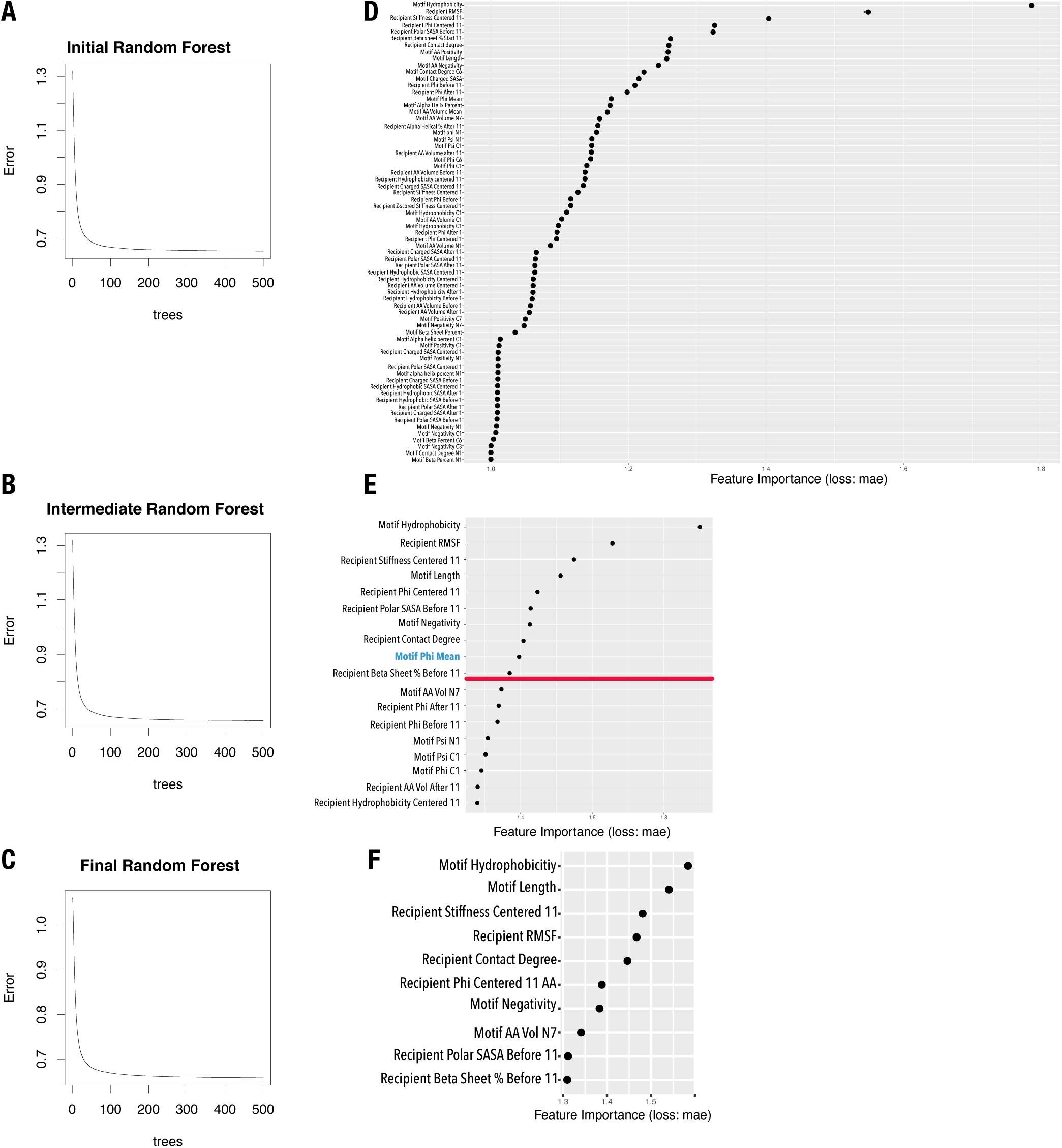
Random Forest model iteration training and property importance. (**A-C**) Error curves with mean squared error plotted against number of trees in the (**A**) initial, (**B**) intermediate, and (**C**) final Random forest models. As more trees are added there is less error. (**D-F**) Bar plots of the importance of features in predicting insertional fitness in the (**D**) initial, (**E**) intermediate, and (**F**) final Random Forest models. In (**E**) the threshold that was used to trim features is marked with a red line. In addition, mean motif phi angle (blue) was removed because it required motifs to have solved structures, which substantially limited the number of motifs we could include. Property importance is based on the mean absolute error (mae) of removing properties from the predictive model. Further details can be found in the *Materials and Methods*.

**Supplemental Figure 6:**
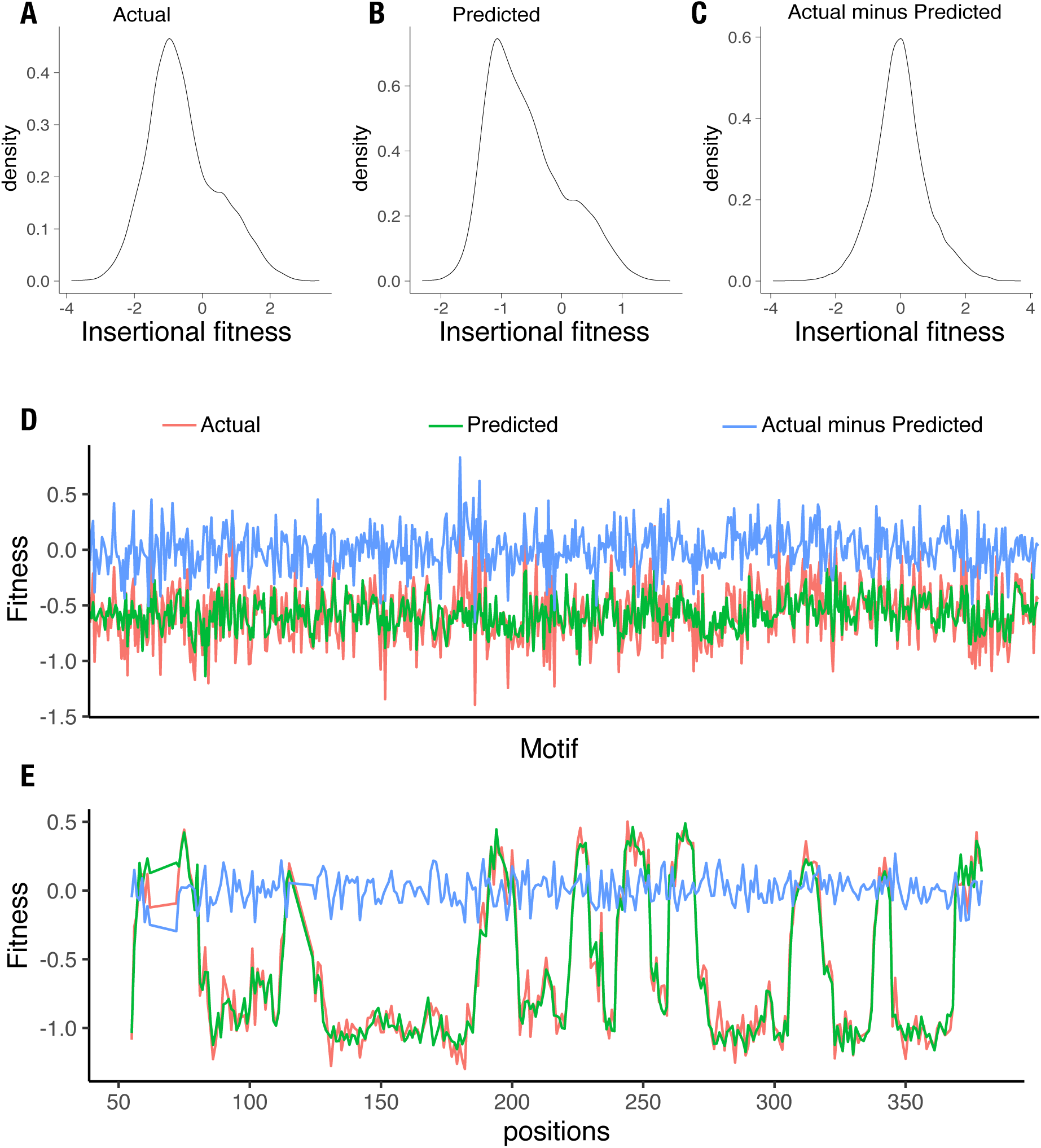
Model performance plots. (**A-C**) Density plots for (**A**) actual, (**B**) predicted, and (**C**) difference between actual and predicted insertional fitness. (**D**) Insertional fitness actual, predicted, and the difference per domain. (**E**) Insertional fitness actual, predicted, and difference per recipient insertion position. All model performance is reported based on data withheld from all random forest training.

**Supplemental Figure 7:**
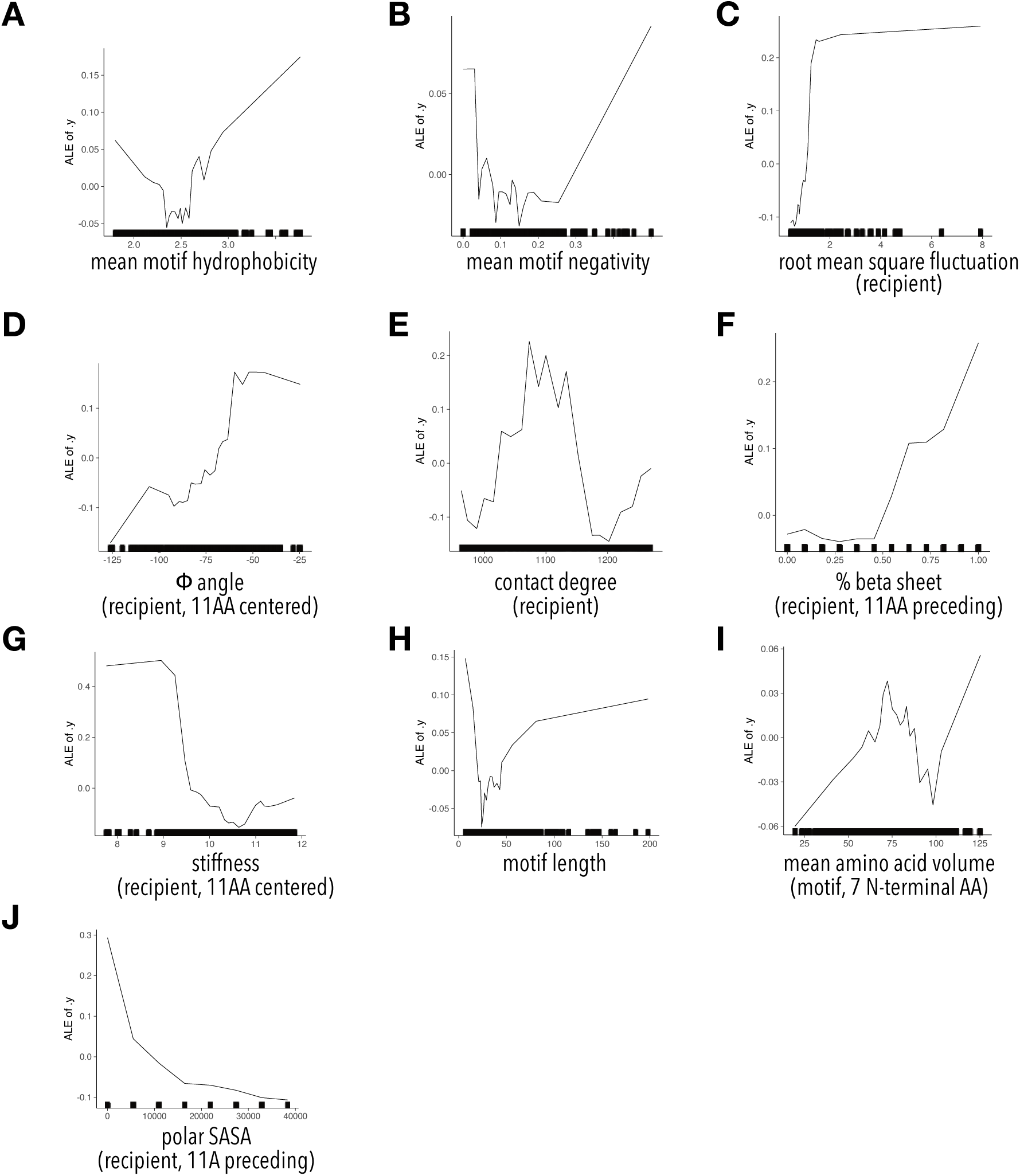
ALE plots for final model properties. Plots of the Accumulated Local Effects (ALE) of properties on the prediction of insertional fitness for (**A**) mean motif hydrophobicity, (**B**) mean motif negativity, (**C**) recipient root mean square fluctuation (based on MD simulation, PDB code 3JYC), (**D**) mean recipient phi angles of 11 AA centered around insertion site, (**E**) recipient contact density, (**F**) mean beta sheet content 11 AA before insertion site (**G**) mean recipient stiffness of 11 AA centered around insertion site, (**H**) motif length, (**I**) mean amino acid volume of the motif’s 7 N terminal AA, and (**J**) polar surface accessible surface area of 11 AA before insertion site.

**Supplemental Figure 8:**
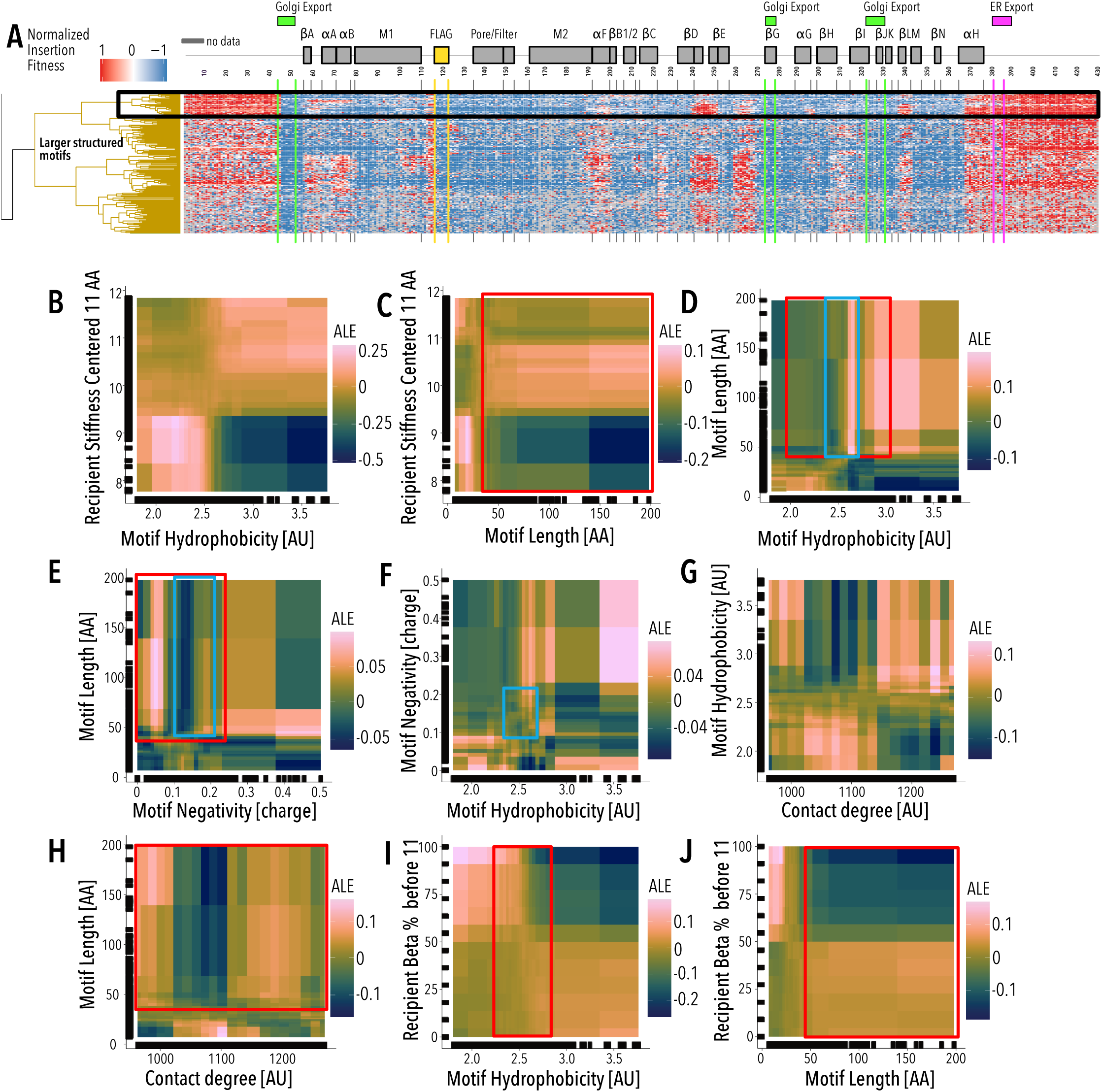
Larger structured motif cluster pairwise ALE Exploration. (**A**) Insertion fitness heatmap of structured motifs inserted into all positions of Kir2.1. Secondary structural elements (grey boxes) for Kir2.1 are shown above, along known Golgi and ER export signals (green and magenta boxes, respectively). Motifs are hierarchically clustered by on a cosine distance metric. The black box indicates a subset of ‘well-structured motifs’ (see Fig. 2F-H). (**B-J**) Pairwise ALE plots investigate how pairwise interactions contribute to prediction of (**B**) recipient stiffness - motif hydrophobicity, (**C**) recipient stiffness - motif length, (**D**) motif hydrophobicity - motif length, (**E**) motif length-motif hydrophobicity, (**F**) motif negativity - motif hydrophobicity, (**G**) motif hydrophobicity-recipient contact degree, (**H**) motif length - recipient contact degree, (**I**) recipient Beta %- motif hydrophobicity, and (**J**) recipient beta%- motif length. Pairwise ALE plots are colored from dark blue to pink with increasing ALE scores. The distribution of larger motifs cluster is boxed in red and the distribution of the well-structured is boxed in blue. Marginal ticks (**B–J**) indicate data point used in model building.

**Supplemental Figure 9:**
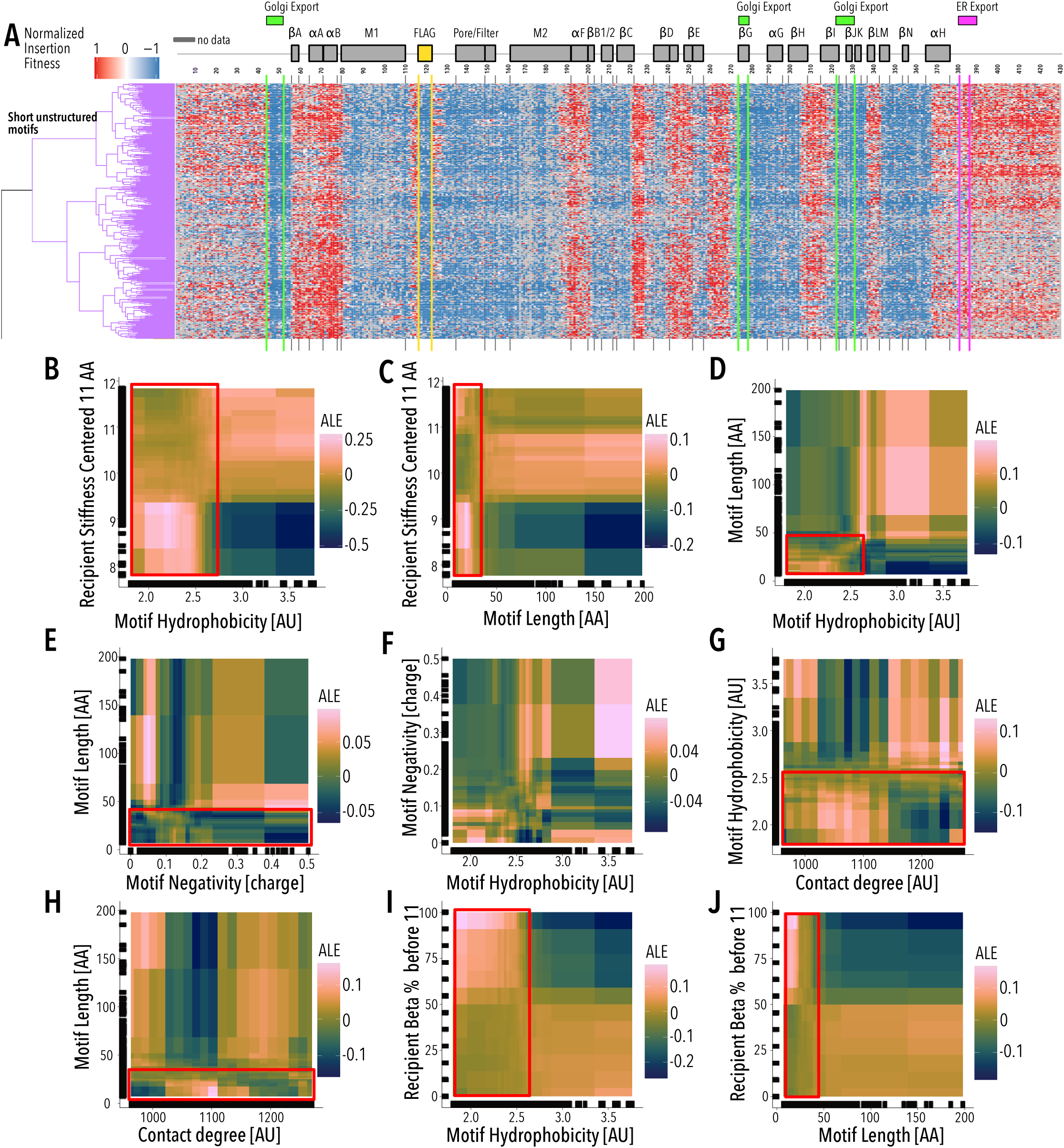
Short unstructured motif cluster pairwise ALE Exploration. (**A**) Insertion fitness heatmap of short unstructured motifs inserted into all positions of Kir2.1. Secondary structural elements (grey boxes) are Kir2.1 are shown above, along known Golgi and ER export signals (green and magenta boxes, respectively). Motifs are hierarchically clustered by on a cosine distance metric. (**B-J**) Pairwise ALE plots investigate how pairwise interactions contribute to prediction of (**B**) recipient stiffness - motif hydrophobicity, (**C**) recipient stiffness - motif length, (**D**) motif hydrophobicity - motif length, (**E**) motif length - motif hydrophobicity, (**F**) motif negativity- motif hydrophobicity, (**G**) motif hydrophobicity - recipient contact degree, (**H**) motif length - recipient contact degree, (**I**) recipient Beta % - motif hydrophobicity, and (**J**) recipient beta % - motif length. Pairwise ALE plots are colored from dark blue to pink with increasing ALE scores. The distributions of hydrophobic motifs cluster are boxed in red. Marginal ticks (**B–J**) indicate data point used in model building.

**Supplemental Figure 10:**
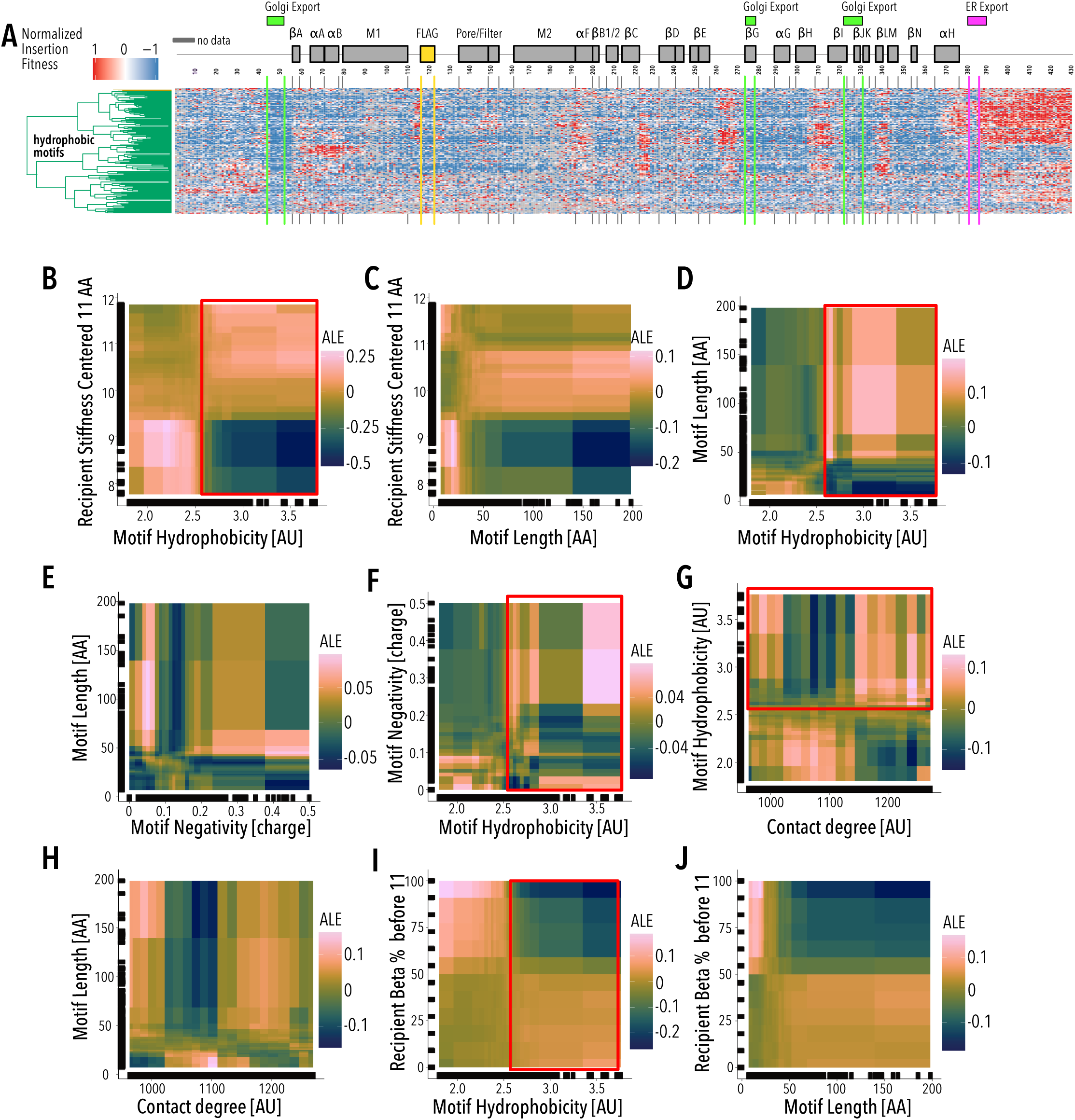
Hydrophobic motif cluster pairwise ALE Exploration. (**A**) Insertion fitness heatmap of hydrophobic motifs inserted into all positions of Kir2.1. Secondary structural elements (grey boxes) for Kir2.1 are shown above, along known Golgi and ER export signals (green and magenta boxes, respectively). Motifs are hierarchically clustered by on a cosine distance metric. (**B-J**) Pairwise ALE plots investigate how pairwise interactions contribute to prediction of (**B**) recipient stiffness-motif hydrophobicity, (**C**) recipient stiffness-motif length, (**D**) motif hydrophobicity-motif length, (**E**) motif length-motif hydrophobicity, (**F**) motif negativity - motif hydrophobicity, (**G**) motif hydrophobicity - recipient contact degree, (**H**) motif length-recipient contact degree, (**H**) recipient Beta % - motif hydrophobicity, and (**J**) recipient beta % - motif length. Pairwise ALE plots are colored from dark blue to pink with increasing ALE scores. The distributions of hydrophobic motifs cluster are boxed in red. Marginal ticks (**B–J**) indicate data point used in model building.

**Supplemental Figure 11:**
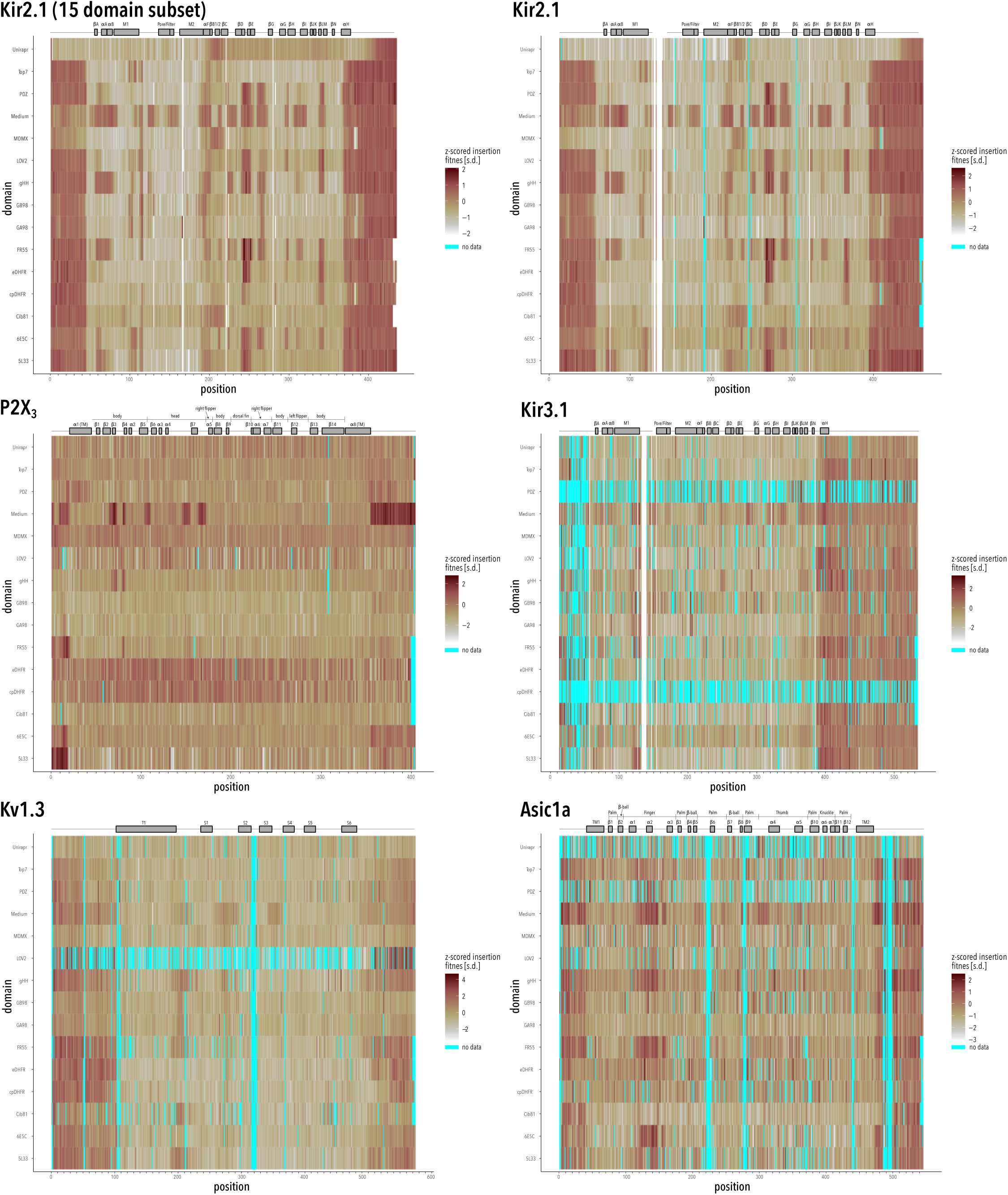
Mean insertion fitness across channels and domains. All datasets are based on at least two biological replicates. Two datasets are shown for Kir2.1 that were collected with different sequencing chemistry. Secondary structure elements (and topological organization; P2X_3_ and Asic1a only) are shown as cartoons.

**Supplemental Figure 12:**
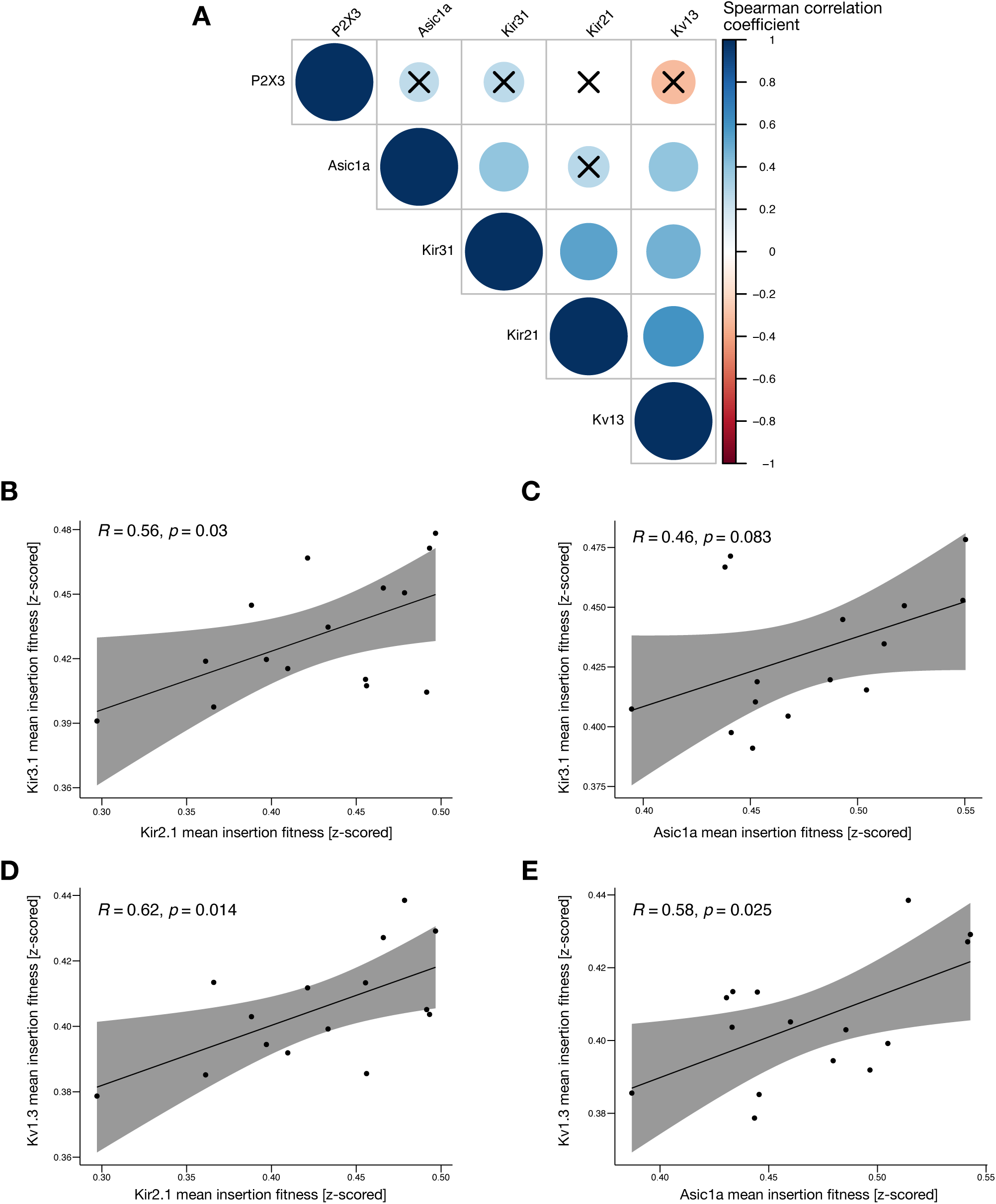
Correlation of domain insertion fitness in different ion channels. (**A**) Spearman correlation of mean insertion fitness (across all channel positions and motifs) between different channel pairs. Crosses indicate coefficient p-values > 0.05 (i.e., not significant). (**B-E**) Scatterplots of mean insertion fitness (across all channel position) for each inserted motifs. The solid black line indicates a linear regression and the grey shaded area indicates a 90% confidence interval. Spearman correlation coefficient and p-value are shown for each channel combination. Overall, correlation of motif effects on insertion fitness is moderate, suggesting a minor role relative to recipient channel properties.

**Supplemental Figure 13:**
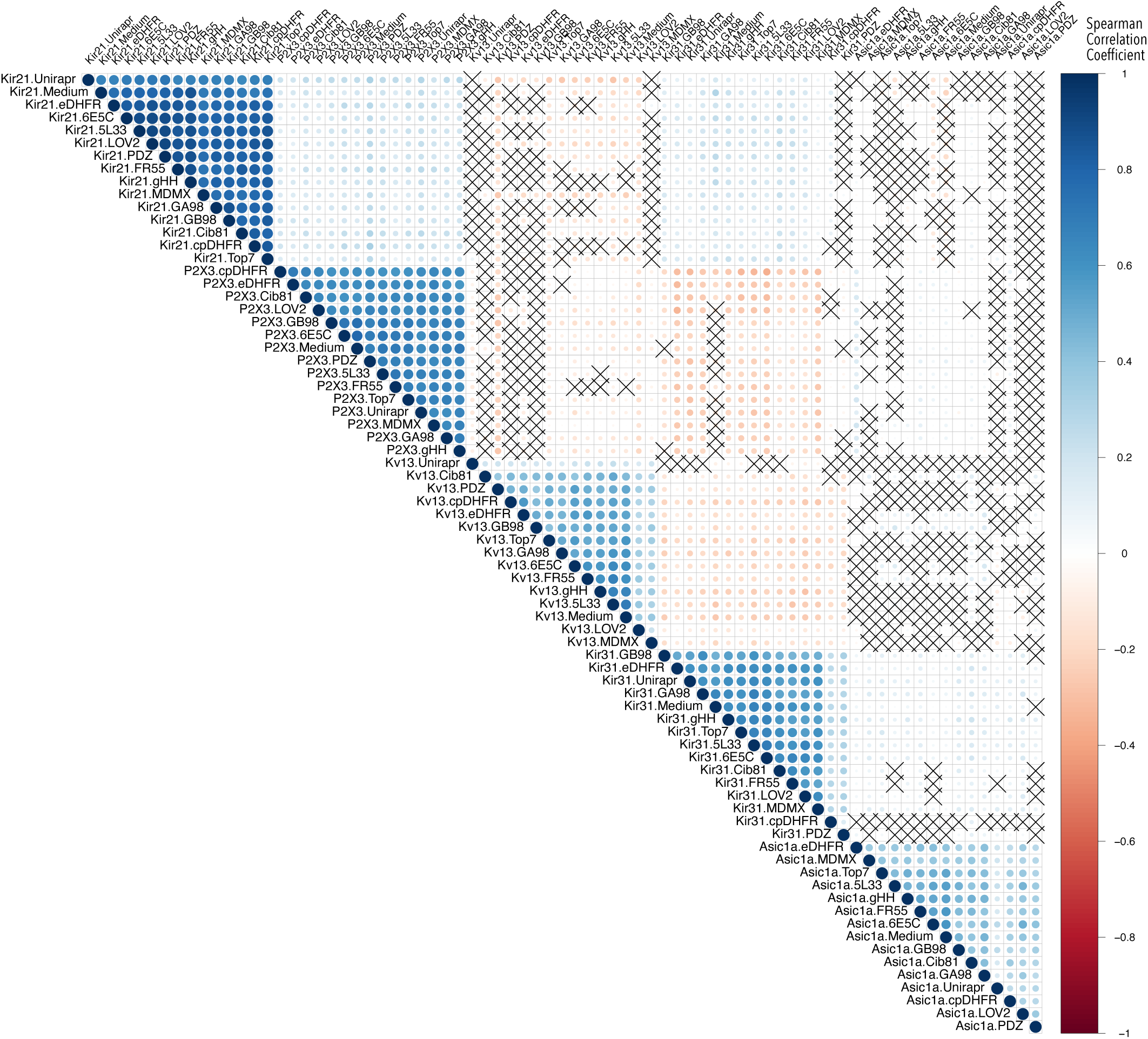
Correlations of insertion fitness for motifs in different channels. Spearman correlation of mean insertion fitness (across all channel position) of a specific motif in a specific channels with all other combinations. Strong correlations of different motifs in the same channel background dominate, suggesting that the recipient properties’ influence on fitness is strong. Crosses indicate coefficient p-values > 0.05 (i.e., not significant).

**Supplemental Figure 14:**
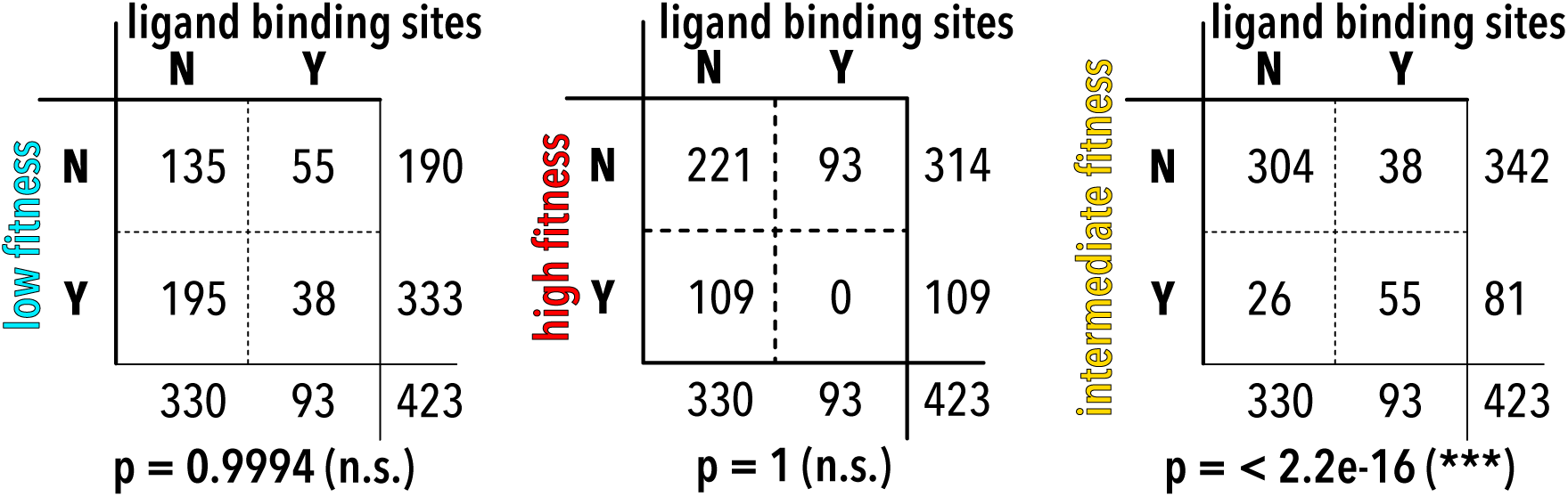
Class / ligand binding sites contigency tables. Independence of inward rectifier ligand binding sites (PIP_2_ – Kir2.1, Kir3.1, Kir6.2, Gβγ – Kir3.1 only, ATP – Kir6.2 only) with respect to different residue classes identified by unbiased clustering of insertion fitness was tested using two-sided Fisher’s Exact tests. Only the intermediate fitness class (colored yellow in Fig. 1D) is enriched for ligand binding sites.

**Supplemental Figure 15:**
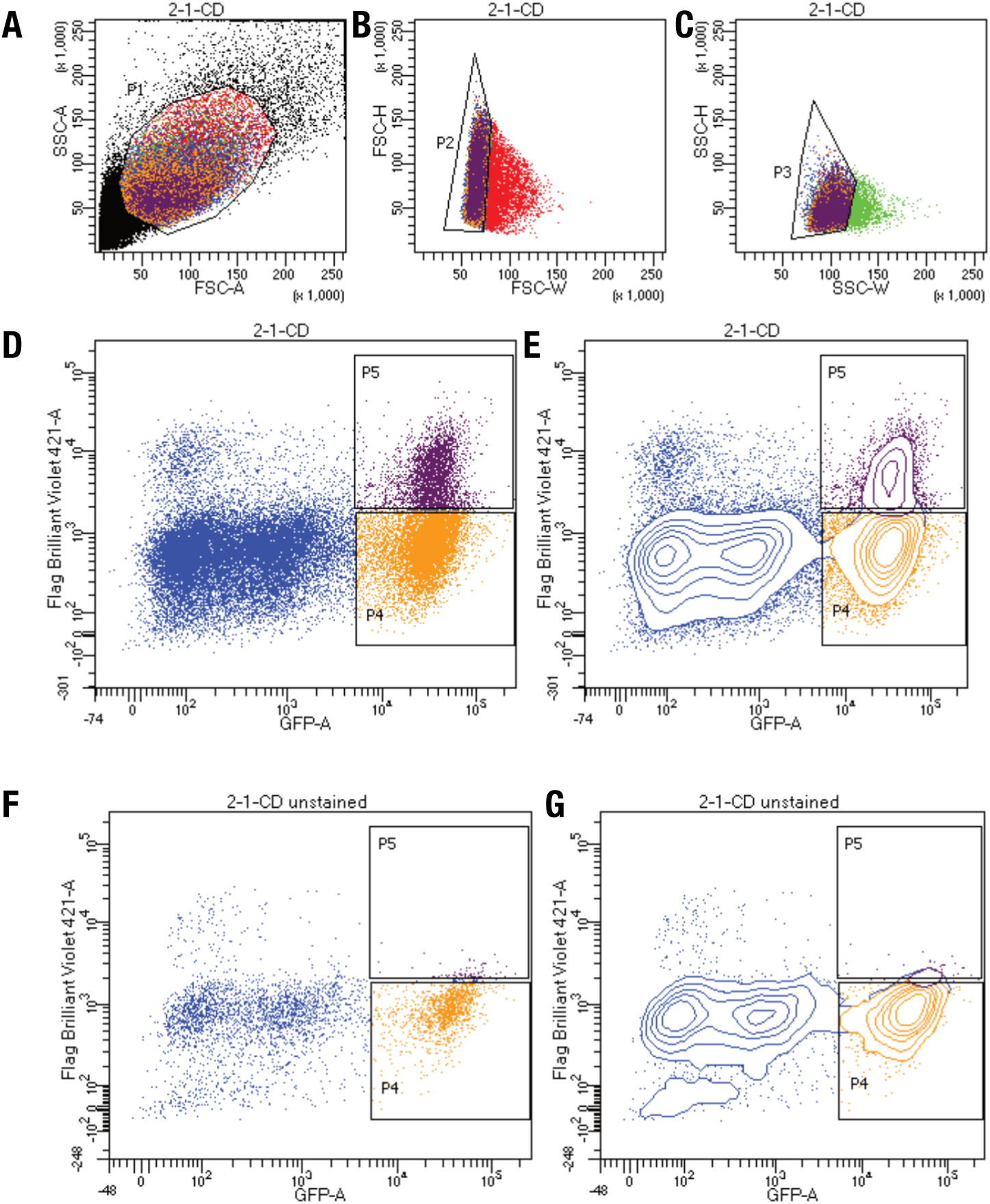
Kir2.1 surface expression assay gating scheme. (**A**) Whole HEK293 cells are gated on side (SSC-A) and forward scattering (FSC-A). (**B-C**) Forward scattering height (SSC-H), forward scattering width (FSC-W), and Side scattering width (SSC-W) are used to gate single cells. (**D-G**) EGFP^high^/Label^low^ and EGFP^high^/Label^high^ populations are gated based (**D-E**) stained and (**F-G**) unstained on EGFP (GFP-A) of Anti-Flag Brilliant Violet-421 fluorescence with (**D,F**) scatterplot and (**E,G**) contour plots shown. Contour plots represent 95% confidence intervals with outliers shown as dots.

**Supplemental Figure 16:**
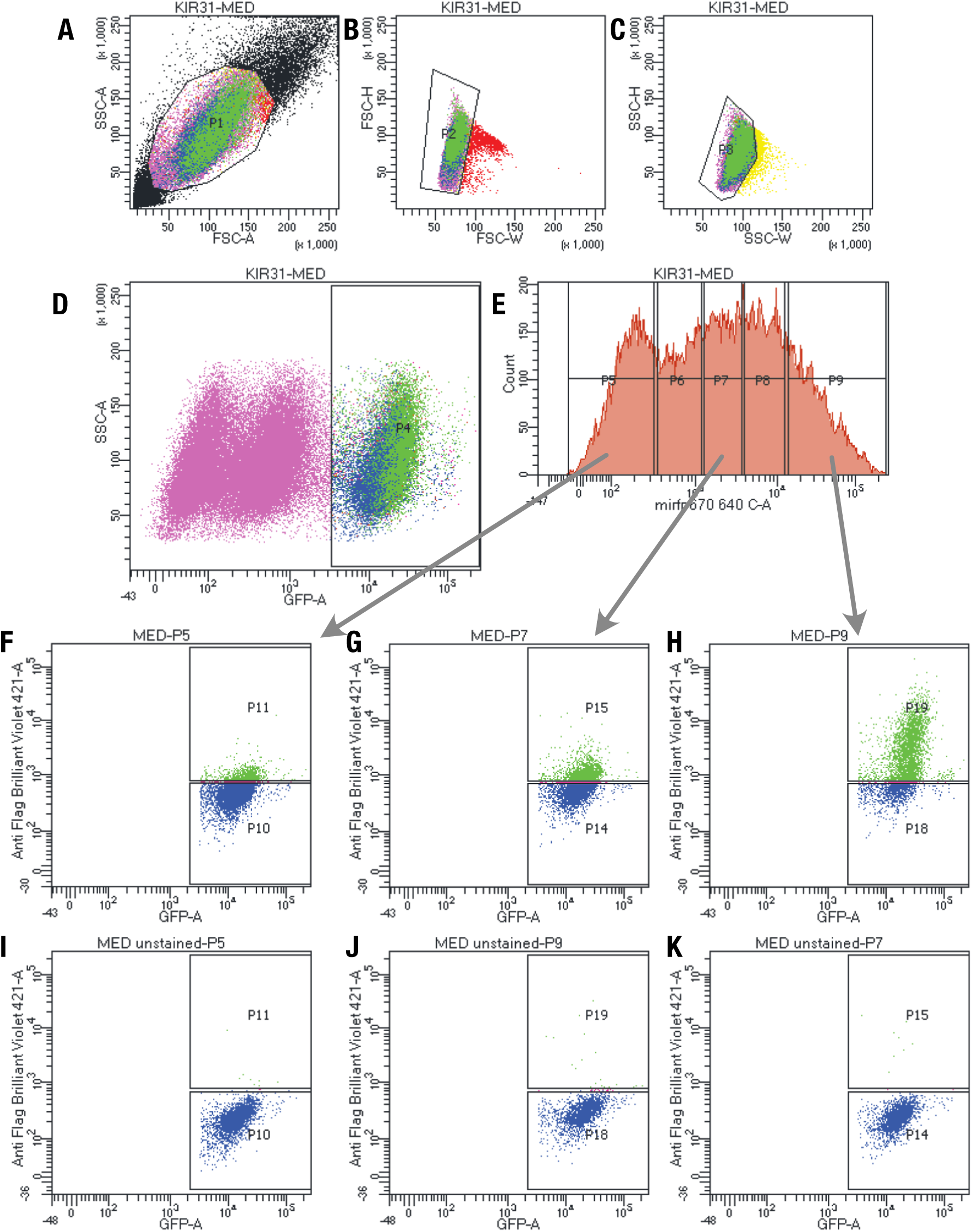
Kir3.1 surface expression assay gating scheme. (**A**) Whole HEK293 cells are gated on side (SSC-A) and forward scattering (FSC-A). (**B-C**) Forward scattering height (SSC-H), forward scattering width (FSC-W), and Side scattering width (SSC-W) are used to gate single cells. (**D**) Cells are gated on EGFP positive cells to isolate successfully recombined libraries. (**E**) Cells are further split into 5 populations to separate out different populations of Kir3.2 co-expressed miRFP670. (**F-K**) EGFP^high^/ Label^low^ and EGFP^high^/ Label^high^ populations are gated based (**F-H**) stained and (**I-K**) unstained on EGFP (GFP-A) of Anti-Flag Brilliant Violet-421 fluorescence. The data from 3 highest levels of miRFP670 were combined and reported as fitness.

**Supplemental Figure 17:**
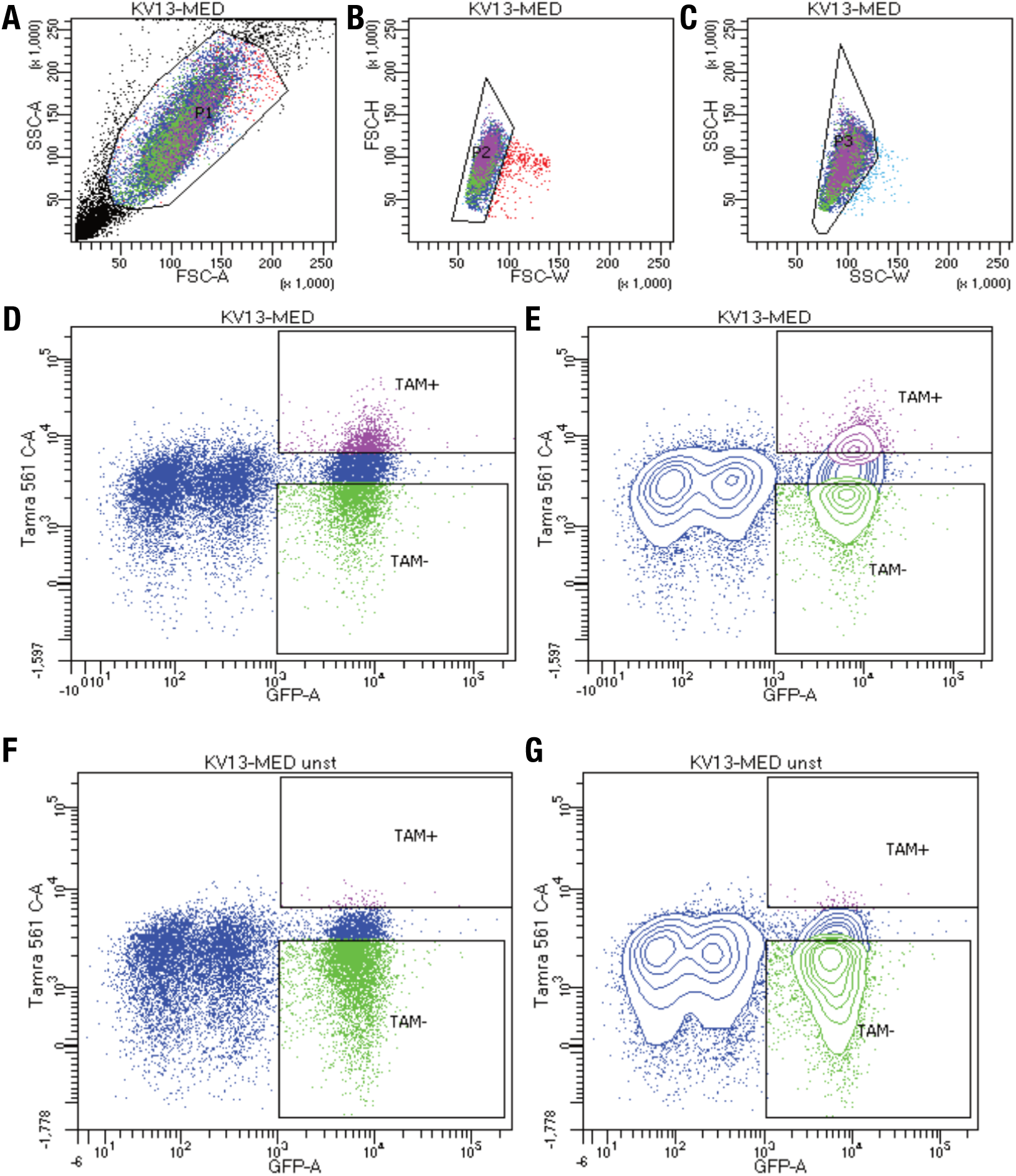
Kv1.3 Surface expression assay gating scheme. (**A**) Whole HEK293 cells are gated on side (SSC-A) and forward scattering (FSC-A). (**B-C**) Forward scattering height (SSC-H), forward scattering width (FSC-W), and Side scattering width (SSC-W). (**D-G**) EGFP^high^/ Label^low^ and EGFP^high^/Label^high^ populations are gated based (**D-E**) stained and (**F-G**) unstained on EGFP (GFP-A) of Kv1.3 specific Agitoxin-Tamra fluorescence with (**D,F**) scatterplot and (**E,G**) contour plots shown. Contour plots represent 95% confidence intervals with outliers shown as dots.

**Supplemental Figure 18:**
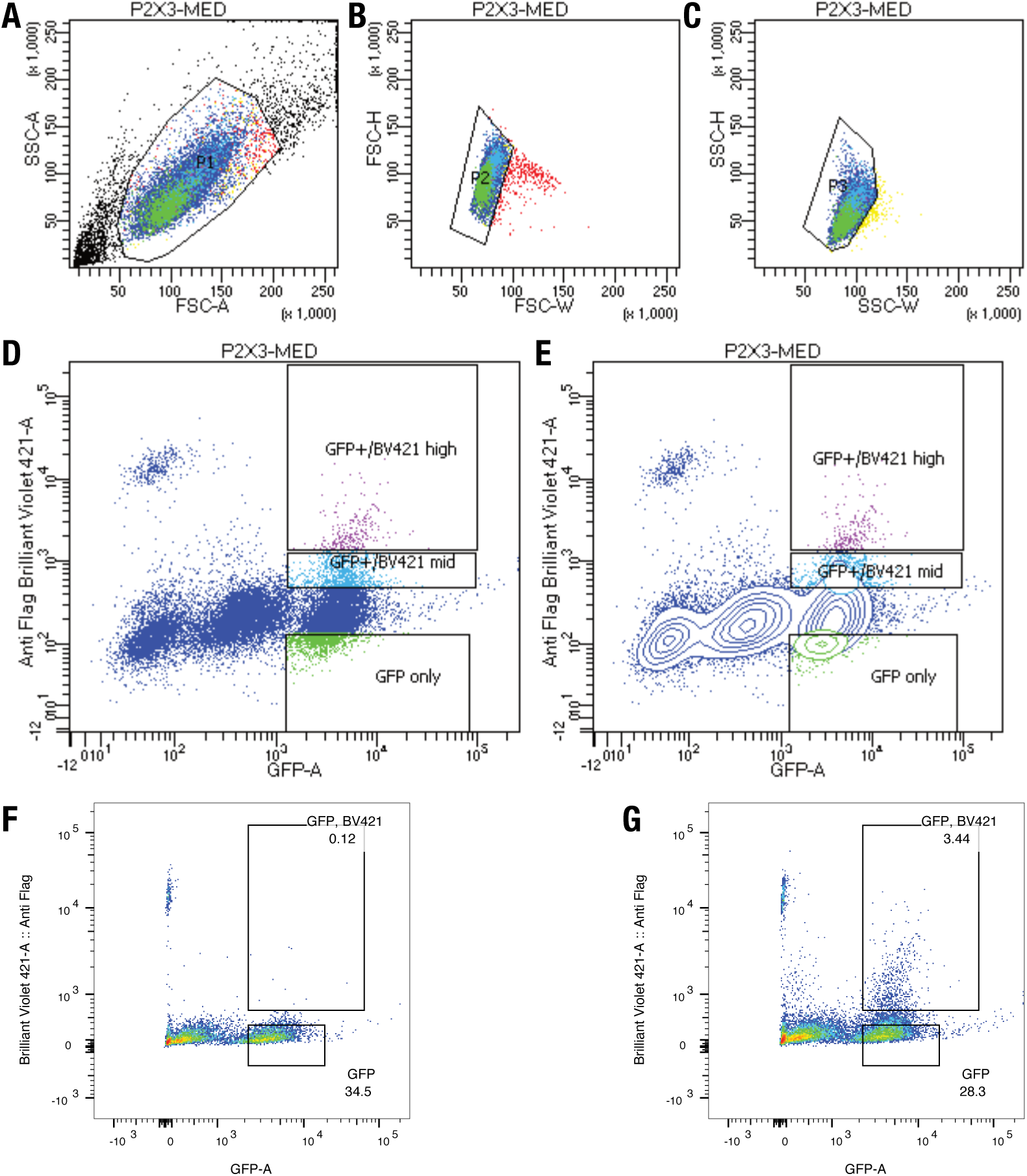
P2X_3_ Surface expression assay gating scheme. (**A**) Whole HEK293 cells are gated on side (SSC-A) and forward scattering (FSC-A). (**B-C**) Forward scattering height (SSC-H), forward scattering width (FSC-W), and Side scattering width (SSC-W) are used to gate single cells. (**D-E**) EGFP^high^/ Label^low^, EGFP ^high^/Label^low^, EGFP^high^/Label^med^ and EGFP^high^/Label^high^ populations are gated based (**F**) stained and (**G**) unstained on EGFP (GFP-A) of Anti-Flag Brilliant Violet-421 fluorescence with (**D**) scatterplot, (**E**) Contour plot, and (**F-G**) pseudo color plots. In post sample collection *Mid* and *High* label populations were combined ratiometrically based on percent populations in corresponding gates. Contour plots represent 95% confidence intervals with outliers shown as dots. Pseudocolor plots represent density of points with a blue-to-red color scale with increasing density.

**Supplemental Figure 19:**
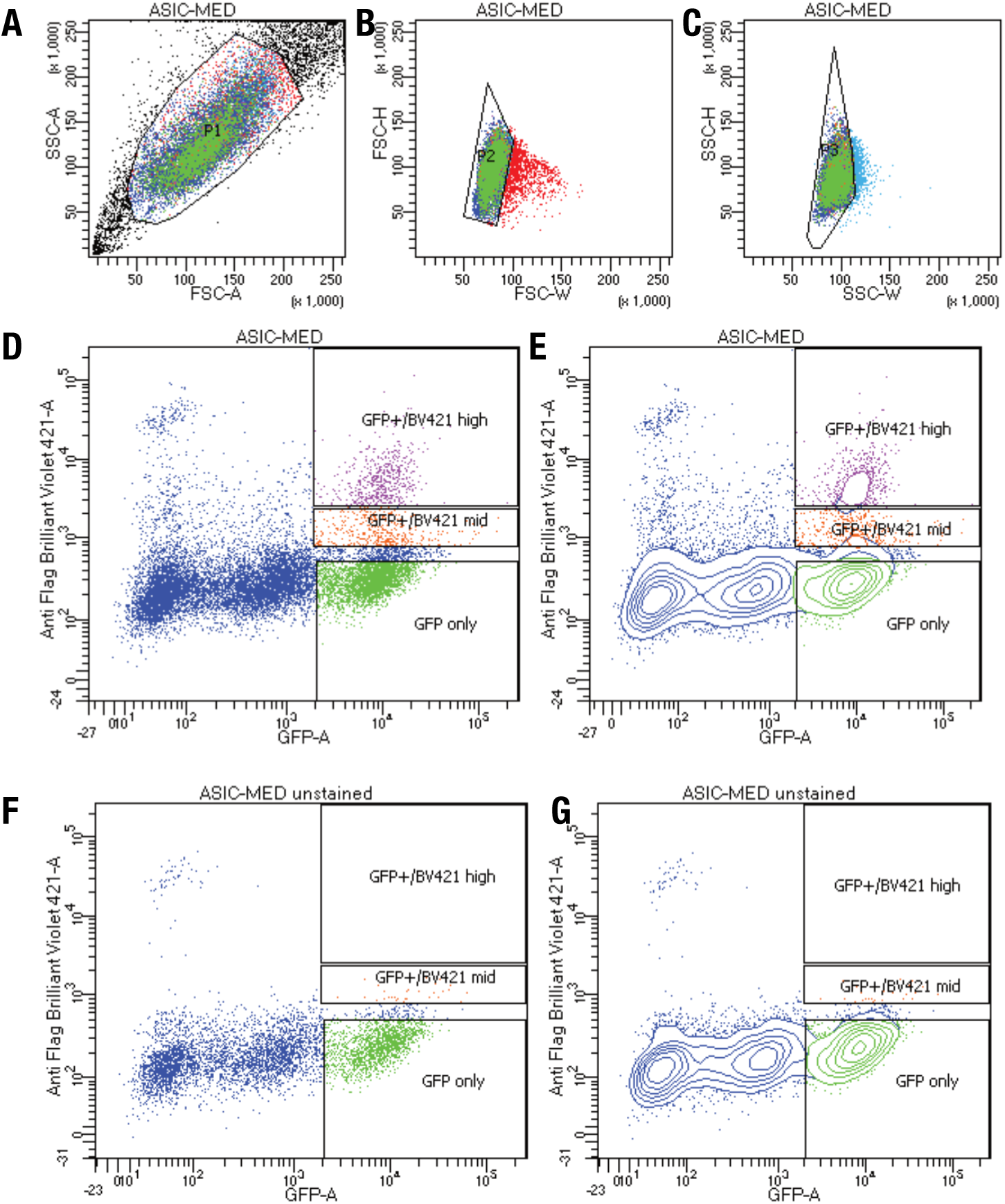
ASIC1a Surface expression assay gating scheme. (**A**) Whole HEK293 cells are gated on side (SSC-A) and forward scattering (FSC-A). (**B-C**) Forward scattering height (SSC-H), forward scattering width (FSC-W), and Side scattering width (SSC-W) are used to gate single cells. (**D-G**) EGFP^high^/ Label^low^ and EGFP^high^/Label^high^ populations are gated based (**D-E**) stained and (**F-G**) unstained on EGFP (GFP-A) of Anti-Flag Brilliant Violet-421 fluorescence with (**D,F**) scatterplot and (**E,G**) contour plots shown. Contour plots represent 95% confidence intervals with outliers as shown as dots.

**Supplemental Figure 20:**
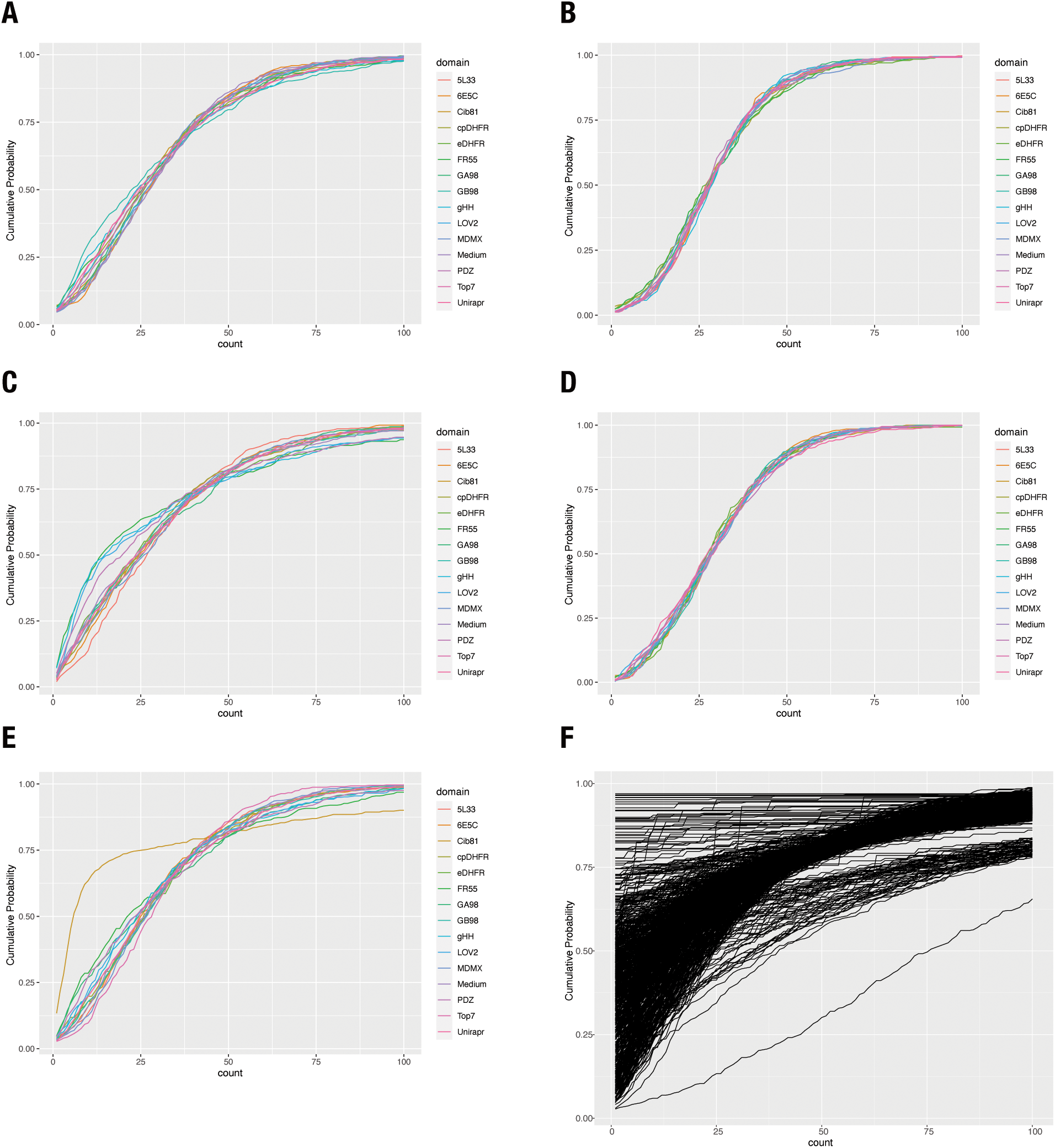
Baseline profiles for each domain and gene combination. (**A-F**) Empirical cumulative distribution plots for ASIC1a (**A**), Kir2.1 (**B**), Kir3.1 (**C**), P2X_3_ (**D**), and Kv1.3 (**E**). (**F**) Large domain set for Kir2.1. Each domain was normalized to have 30x coverage before calculating empirical cumulative distribution function. Plots show cumulative probability for each count threshold from 1 to 100. This indicates distribution of insertions in a given gene with distributions shifted to the right being more evenly distributed.

**Supplemental Table 1:**
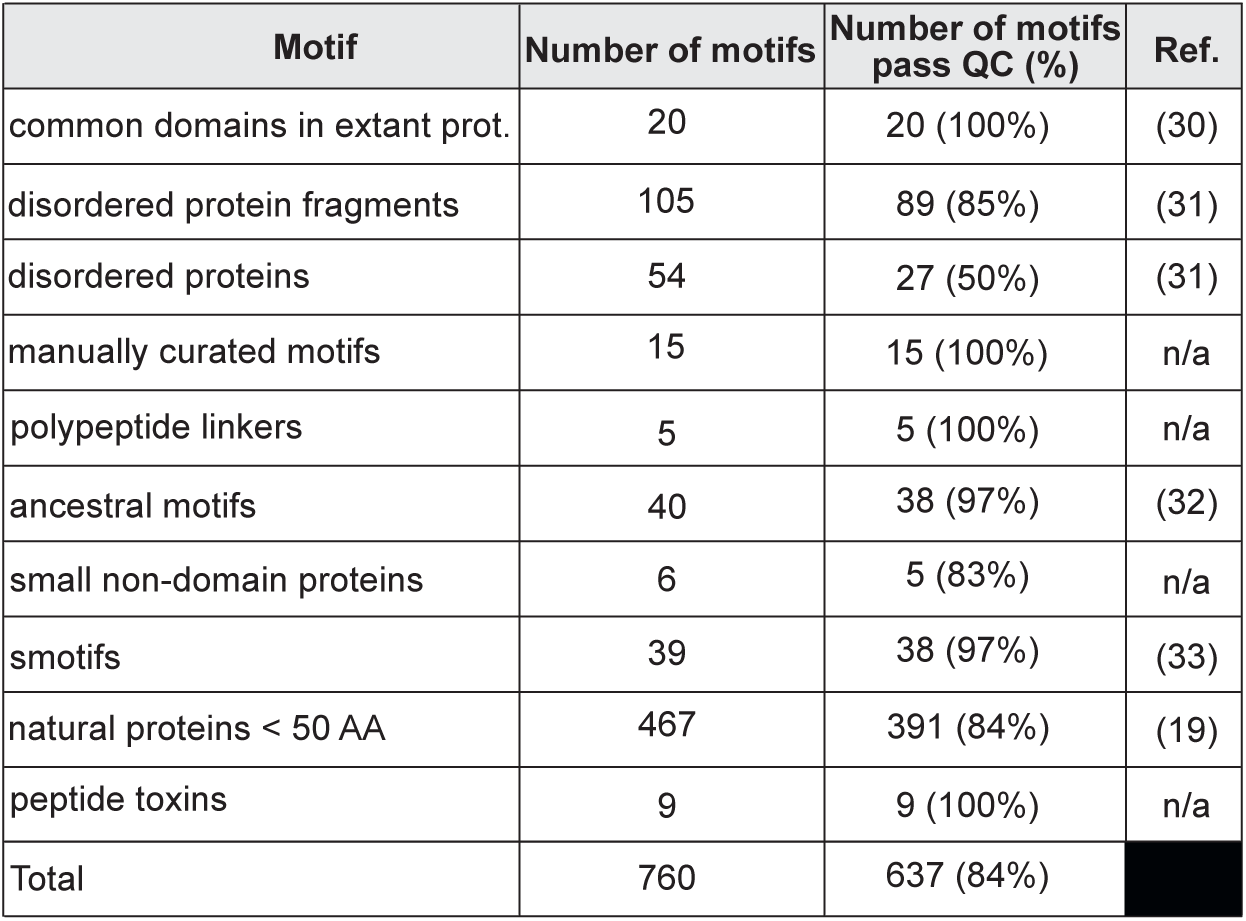
Motif group statistics for Kir2.1 760 motif dataset. Number of motifs, number of motifs passing QC threshold, and sources. Motifs pass QC if they contain statistically significant data in greater than 80% of insertion positions and are included in further analysis and model building.

**Supplemental Table 2:** Inserted motif properties. This table only contains means and standard deviations of the insertion position properties. All additional sliding window recipient properties are provided in a supplemental .csv file. Å refers to Angstroms, AA refers to amino acids, Da refers to Daltons, and AU to arbitrary units.

| Motif property | Abbreviation | Mean +/- SD | Reference |
| --- | --- | --- | --- |
| Motif Length [AA] | Motif_length | 37.2 +/- 22.2 | n/a |
| Phi Mean [degrees] | d_phi_mean | -68.0 +/- 12.4 | (45) |
| Psi Mean [degrees] | d_psi_mean | -0.49 +/- 35.9 | (45) |
| Radius of Gyration [Å] | d_gyradius | 12.3 +/- 3.3 | (45) |
| NC distance [Å] | d_nc_dist | 12.3 +/- 3.3 | (45) |
| Distance of N term to center of mass [Å] | d_center_n_dist | 23.8 +/- 12.8 | (45) |
| Distance of C term to center of mass [Å] | d_center_c_dist | 23.0 +/- 11.8 | (45) |
| Contact degree [AU] | d_contact_degree | 450 +/- 287 | (39) |
| Contact order [AU] | d_contact_order | 0.41 +/- 0.038 | (39) |
| Long contact degree [AU] | d_long_degree | 7.99 +/- 9.12 | (39) |
| Secondary Structure (%) | d_sspercent | 60.0 +/- 25.0 | (39) |
| Alpha helical [%] | d_alpha_percent | 53.9 +/- 31.3 | (39) |
| Beta sheet [%] | d_beta_percent | 6.1 +/- 13.6 | (39) |
| Buried nonpolar surface area [Å<sup>2</sup>] | d_npsa | 2100 +/- 1990 | (39) |
| Charged solvent accessible surface area [Å<sup>2</sup>] | d_charged_mean | 39,600 +/- 53,700 | (39) |
| Polar solvent accessible surface area [Å<sup>2</sup>] | d_polar_mean | 40,710 +/- 56,000 | (39) |
| Hydrophobic solvent accessible surface area [Å<sup>2</sup>] | d_hydrophob_mean | 69,000 +/- 88,000 | (39) |
| Root mean squared deviation between conformers | d_rmsd | 2.98 +/- 2.25 | (45) |
| Stiffness [AU] | d_stiffness_mean | -7.62E-18 +/- 1.08 E-15 | (38) |
| Mean AA Molecular Weight [Da] | d_AA_MW_mean | 130. +/- 7.49 | (46) |
| Mean AA Surface area [Å<sup>2</sup>] | d_AA_SA_mean | 158 +/- 16 | (46) |
| Mean AA Alpha helical propensity [AU] | d_AA_alphahel_mean | 1.04 +/- 0.07 | (46) |
| Mean AA Beta sheet propensity [AU] | d_AA_betashe_mean | 0.99 +/- 0.07 | (46) |
| Mean AA Buried accessibility ratio propensity [AU] | d_AA_bur_acc_ratio_mean | 1.25 +/- 0.29 | (46) |
| Mean AA flexibility [AU] | d_AA_flex_mean | 0.44 +/- 0.02 | (46) |
| Mean AA hydropathy [AU] | d_AA_hydropath_mean | -0.43 +/- 0.84 | (46) |
| Mean AA hydrophobicity [AU] | d_AA_hydrophob_mean | 2.5 +/- 0.26 | (46) |
| Mean AA negative charge | d_AA_negat_mean | 0.117 +/- 0.079 | (46) |
| Mean AA pka | d_AA_pka_mean | 4.28 +/- 0.28 | (46) |
| Mean AA polarity [AU] | d_AA_polar_mean | 8.6 +/- 0.7 | (46) |
| Mean AA positive charge | d_AA_posit_mean | 0.17 +/- 0.09 | (46) |
| Mean AA reverse turn propensity [AU] | d_AA_rev_turn_mean | 0.97 +/- 0.11 | (46) |
| Mean AA volume [Å<sup>3</sup>] | d_AA_vol_mean | 79.1 +/- 10.7 | (46) |
| Length of structures [AA] | d_size | 36.6 +/- 20.1 | (39) |

**Supplemental Table 3:** Recipient insertion position properties. This table only contains means and standard deviations of the insertion position properties. All additional sliding window recipient properties are provided in a supplemental .csv file. Å refers to Angstroms, AA refers to amino acids, Da refers to daltons, and AU to arbitrary units.

| Recipient insertion position property | Abbreviation | Mean +/- SD | Reference |
| --- | --- | --- | --- |
| MD Root mean square fluctuation 3SPI (AU) | rmsf_3spi | 0.96 +/- 0.70 | n/a |
| MD Root mean square fluctuation 3JYC(AU) | rmsf_3jyc | 1.14 +/- 0.85 | n/a |
| Phi (Degrees) | phi | -75.6 +/- 57.7 | (45) |
| Psi (Degrees) | psi | 41.3 +/- 88.4 | (45) |
| Contact degree (AU) | cdegree | 1116.5 +/- 92.0 | (39) |
| Contact order (AU) | corder | 0.439 +/- 0.036 | (39) |
| Long contact degree (AU) | longdegree | 0.863 +/- 0.072 | (39) |
| Secondary Structure (percentage) | ss | 0.60 +/- 0.49 | (39) |
| Alpha helix (percentage) | alpha | 0.33 +/- 47 | (39) |
| Beta sheet (percentage) | beta | 0.27 +/- 0.44 | (39) |
| Buried nonpolar surface area (Å<sup>2</sup>) | npsa | -12.2 +/- 144.4 | (39) |
| Charged solvent accessible surface area (Å<sup>2</sup>) | charged_sasa | 13,069 +/- 24,866 | (39) |
| Polar solvent accessible surface area (Å<sup>2</sup>) | polar_sasa | 16,100 +/- 26,697 | (39) |
| Normal Mode based Stiffness (AU) | stiffness | 10.33 +/- 1.12 | (38) |
| AA Surface area (Å<sup>2</sup>) | AA_SA | 159.5 +/- 57.9 | (46) |
| AA Buried accessibility ratio propensity (AU) | AA_bur_acc_ratio | 1.41 +/- 1.17 | (46) |
| AA Alpha helical propensity (AU) | AA_alpha_hel | 1.03 +/- 0.25 | (46) |
| AA Beta sheet propensity (AU) | AA_beta_she | 1.02 +/- 0.26 | (46) |
| AA reverse turn propensity (AU) | AA_rev_turn | 0.94 +/- 0.38 | (46) |
| AA volume (Å<sup>3</sup>) | AA_vol | 79.7 +/- 39.1 | (46) |
| AA flexibility (AU) | AA_flex | 0.438 +/- 0.075 | (46) |
| AA Buried accessibility ratio propensity (AU) | AA_bur_vol | 146 +/- 39 | (46) |
| AA Molecular weight (Da) | AA_MW | 131 +/- 27 | (46) |
| AA positive charge | AA_posit | 0.133 +/- 0.340 | (46) |
| AA negative charge | AA_negat | 0.140 +/- 0.35 | (46) |
| AA pka | AA_pka | 4.33 +/- 1.02 | (46) |
| AA polarity (AU) | AA_polar | 8.43 +/- 2.72 | (46) |
| AA hydropathy (AU) | AA_hydropath | -0.133 +/- 3.138 | (46) |
| AA Hydrophobicity (AU) | AA_hydrophob | 2.62 +/- 1.02 | (46) |

**Supplemental Table 4:** Random forest parameters. Despite substantially reducing the number of properties, model performance based on variance explained and mean squared residuals are not significantly impacted.

| Random Forest | Variance explained (%) | Mean square residuals | Recipient properties (#) | Motif properties (#) | Total properties (#) |
| --- | --- | --- | --- | --- | --- |
| Initial | 39.89 | 0.652 | 37 | 32 | 69 |
| Intermediate | 39.44 | 0.657 | 10 | 8 | 18 |
| Final | 38.69 | 0.658 | 6 | 4 | 10 |

**Supplemental table 5:** Smaller set of 15 motifs.

| Motif | Length (AA) | Natural or designed | ref |
| --- | --- | --- | --- |
| AGSAGSA | 7 | Designed | n/a |
| Syntrophin PDZ | 86 | Natural | (48) |
| Cib81 | 81 | Natural | (49) |
| e. coli cpDHFR | 164 | Modified | (50) |
| e. coli DHFR | 164 | Natural | (51) |
| FR55 | 82 | Designed | n/a |
| GA98 | 56 | Designed | (52) |
| GB98 | 56 | Designed | (52) |
| ghhh06 | 43 | Designed | (53) |
| Unirapr | 198 | Designed | (54) |
| asLOV2 | 143 | Natural | (55) |
| MDMX | 103 | Natural | (56) |
| Top7 | 99 | Designed | (57) |
| 5L33 | 108 | Designed | (58) |
| 6E5C | 73 | Designed | (59) |

**Supplemental Table 6:** Read count statistics.

| Pool Name | Gene | Sample | Replicate | Total Reads in Pool | Mean Quality | Aligned Reads | Number of Domains | Number of Positions | Number of Variants | Coverage [x-fold] |
| --- | --- | --- | --- | --- | --- | --- | --- | --- | --- | --- |
| base_1-16 | Kir2.1 | Baseline | 1 | 81,371,890 | 30.85 | 7482860 | 560 | 436 | 244160 | 30.6 |
| base_17-21 | Kir2.1 | Baseline | 1 | 24,346,753 | 30.92 | 2097244 | 175 | 436 | 76300 | 27.5 |
| base_CD | Kir2.1 | Baseline | 1 | 4,304,294 | 31.17 | 341611 | 45 | 436 | 19620 | 17.4 |
| dp_1_1-16 | Kir2.1 | Surface expression | 1 | 79,873,354 | 30.93 | 7767341 | 560 | 436 | 244160 | 31.8 |
| dp_1_17-21 | Kir2.1 | Surface expression | 1 | 19,242,372 | 30.73 | 1511063 | 175 | 436 | 76300 | 19.8 |
| dp_1_CD | Kir2.1 | Surface expression | 1 | 2,847,220 | 29.28 | 124711 | 45 | 436 | 19620 | 6.4 |
| dp_2_11-12X15 | Kir2.1 | Surface expression | 2 | 9,683,945 | 30.45 | 764060 | 105 | 436 | 45780 | 16.7 |
| dp_2_13-14X18 | Kir2.1 | Surface expression | 2 | 10,853,687 | 31.1 | 311539 | 105 | 436 | 45780 | 6.8 |
| dp_2_19-21X16-17 | Kir2.1 | Surface expression | 2 | 19,264,855 | 29.38 | 1289393 | 175 | 436 | 76300 | 16.9 |
| dp_2_3-10 | Kir2.1 | Surface expression | 2 | 35,490,287 | 30.73 | 2762399 | 280 | 436 | 122080 | 22.6 |
| dp_2_CD | Kir2.1 | Surface expression | 2 | 3,203,291 | 31.36 | 211588 | 45 | 436 | 19620 | 10.8 |
| gfp_1_1-16 | Kir2.1 | No surface expression | 1 | 77,668,001 | 30.82 | 7768024 | 560 | 436 | 244160 | 31.8 |
| gfp_1_17-21 | Kir2.1 | No surface expression | 1 | 27,686,561 | 30.69 | 2270580 | 175 | 436 | 76300 | 29.8 |
| gfp_1_CD | Kir2.1 | No surface expression | 1 | 4,408,355 | 31.78 | 382483 | 45 | 436 | 19620 | 19.5 |
| gfp_2_11-12X15 | Kir2.1 | No surface expression | 2 | 9,433,539 | 31.35 | 937905 | 105 | 436 | 45780 | 20.5 |
| gfp_2_13-14X18 | Kir2.1 | No surface expression | 2 | 9,369,676 | 31.12 | 306212 | 105 | 436 | 45780 | 6.7 |
| gfp_2_19-21X16-17 | Kir2.1 | No surface expression | 2 | 21,731,747 | 30.28 | 2014803 | 175 | 436 | 76300 | 26.4 |
| gfp_2_3-10 | Kir2.1 | No surface expression | 2 | 37,722,582 | 30.96 | 3698554 | 280 | 436 | 122080 | 30.3 |
| gfp_2_CD | Kir2.1 | No surface expression | 2 | 5,126,808 | 30.53 | 415326 | 45 | 436 | 19620 | 21.2 |
|  | Asic1a | High surface expression | 1 |  |  | 330452 | 15 | 546 | 8190 | 40.3 |
|  | Asic1a | Low surface expression | 1 |  |  | 477966 | 15 | 546 | 8190 | 58.4 |
|  | Asic1a | No surface expression | 1 |  |  | 661693 | 15 | 546 | 8190 | 80.8 |
|  | Asic1a | High surface expression | 2 |  |  | 661693 | 15 | 546 | 8190 | 80.8 |
|  | Asic1a | Low surface expression | 2 |  |  | 477966 | 15 | 546 | 8190 | 58.4 |
|  | Asic1a | No surface expression | 2 |  |  | 330452 | 15 | 546 | 8190 | 40.3 |
|  | P2X4 | High surface expression | 1 |  |  | 242095 | 15 | 396 | 5940 | 40.8 |
|  | P2X4 | Low surface expression | 1 |  |  | 609143 | 15 | 396 | 5940 | 102.5 |
|  | P2X4 | No surface expression | 1 |  |  | 464622 | 15 | 396 | 5940 | 78.2 |
|  | P2X4 | High surface expression | 2 |  |  | 328645 | 15 | 396 | 5940 | 55.3 |
|  | P2X4 | Low surface expression | 2 |  |  | 399762 | 15 | 396 | 5940 | 67.3 |
|  | P2X4 | No surface expression | 2 |  |  | 687217 | 15 | 396 | 5940 | 115.7 |
|  | P2X3 | High surface expression | 1 |  |  | 249510 | 15 | 405 | 6075 | 41.1 |
|  | P2X3 | Low surface expression | 1 |  |  | 580254 | 15 | 405 | 6075 | 95.5 |
|  | P2X3 | No surface expression | 1 |  |  | 881800 | 15 | 405 | 6075 | 145.2 |
|  | P2X3 | High surface expression | 2 |  |  | 379725 | 15 | 405 | 6075 | 62.5 |
|  | P2X3 | Low surface expression | 2 |  |  | 489186 | 15 | 405 | 6075 | 80.5 |
|  | P2X3 | No surface expression | 2 |  |  | 918149 | 15 | 405 | 6075 | 151.1 |
|  | Asic2a | High surface expression | 1 |  |  | 455576 | 15 | 520 | 7800 | 58.4 |
|  | Asic2a | Low surface expression | 1 |  |  | 688644 | 15 | 520 | 7800 | 88.3 |
|  | Asic2a | No surface expression | 1 |  |  | 848345 | 15 | 520 | 7800 | 108.8 |
|  | Kv1.3 | Surface expression | 1 |  |  | 434511 | 15 | 575 | 8625 | 50.4 |
|  | Kv1.3 | No surface expression | 1 |  |  | 744064 | 15 | 575 | 8625 | 86.3 |
|  | Kv1.3 | Surface expression | 2 |  |  | 798555 | 15 | 575 | 8625 | 92.6 |
|  | Kv1.3 | No surface expression | 2 |  |  | 810399 | 15 | 575 | 8625 | 94.0 |
|  | Kir3.2 | Surface expression | 1 |  |  | 829744 | 15 | 425 | 6375 | 130.2 |
|  | Kir3.2 | No surface expression | 1 |  |  | 766766 | 15 | 425 | 6375 | 120.3 |
|  | Kir3.2 | Surface expression | 2 |  |  | 988708 | 15 | 425 | 6375 | 155.1 |
|  | Kir3.2 | No surface expression | 2 |  |  | 1025554 | 15 | 425 | 6375 | 160.9 |
|  | Kir2.1 | High surface expression | 1 |  |  | 324519 | 15 | 436 | 6540 | 49.6 |
|  | Kir2.1 | Low surface expression | 1 |  |  | 657683 | 15 | 436 | 6540 | 100.6 |
|  | Kir2.1 | No surface expression | 1 |  |  | 771084 | 15 | 436 | 6540 | 117.9 |
|  | Kir2.1 | High surface expression | 2 |  |  | 613817 | 15 | 436 | 6540 | 93.9 |
|  | Kir2.1 | Low surface expression | 2 |  |  | 521489 | 15 | 436 | 6540 | 79.7 |
|  | Kir2.1 | No surface expression | 2 |  |  | 1110887 | 15 | 436 | 6540 | 169.9 |
|  | Kir6.2 | Surface expression | 1 |  |  | 1230567 | 15 | 410 | 6150 | 200.1 |
|  | Kir6.2 | No surface expression | 1 |  |  | 1377532 | 15 | 410 | 6150 | 224.0 |
|  | Kir6.2 | Surface expression | 2 |  |  | 710829 | 15 | 410 | 6150 | 115.6 |
|  | Kir6.2 | No surface expression | 2 |  |  | 1066743 | 15 | 410 | 6150 | 173.5 |
|  | Kir3.1 | No surface expression | 1 |  |  | 1007295 | 15 | 509 | 7635 | 131.9 |
|  | Kir3.1 | Surface expression | 1 |  |  | 770673 | 15 | 509 | 7635 | 100.9 |
|  | Kir3.1 | No surface expression | 2 |  |  | 764153 | 15 | 509 | 7635 | 100.1 |
|  | Kir3.1 | Surface expression | 2 |  |  | 801705 | 15 | 509 | 7635 | 105.0 |

